# Targeting CD74-positive macrophages improves neoadjuvant therapy in cervical cancer as revealed by single-cell transcriptomics analysis

**DOI:** 10.1101/2023.11.09.566505

**Authors:** Zixiang Wang, Bingyu Wang, Yuan Feng, Jinwen Ye, Zhonghao Mao, Teng Zhang, Meining Xu, Wenjing Zhang, Xinlin Jiao, Youzhong Zhang, Baoxia Cui

**Affiliations:** Department of Obstetrics and Gynecology, Qilu Hospital of Shandong University, Jinan, 250012, P. R. China

**Author notes:** These authors contributed equally: Zixiang Wang, Bingyu Wang. **Correspondence:** Dr. Baoxia Cui, Department of Obstetrics and Gynecology, Qilu Hospital of Shandong University.

**Keywords:** Single-cell RNA sequencing, Cervical cancer, NACT, Tumor microenvironment, Macrophage, CD74

## Abstract

Uterine cervical cancer (UCC) has the second-highest mortality rate among malignant tumors of the female reproductive system. Immune checkpoint inhibitors (ICIs) have emerged as a promising therapeutic strategy for cervical cancer. However, the efficacy of PD-1 (programmed cell death protein 1) blockade combined with neoadjuvant chemotherapy (NACT) and the ensuing alterations within the tumor microenvironment need further investigation. We conducted single-cell RNA sequencing using 46,950 cells from nine sequential human cervical cancer tissues representing different stages of NACT and PD-1 blockade combination therapy. We delineated the trajectory of cervical epithelial cells and unveiled crucial factors involved in the combination therapy. Cell communication analysis revealed the inferred interaction strength decreased between T cell-cancer cells, while the communication between macrophage-cancer cells intensified after NACT therapy. We verified that macrophages are necessary for PD-1 blockade combination to exert antitumor effects in vitro. Detailed analysis unraveled the CD74-positive macrophages frequently interacted with the immunoreactive Epi 3 subgroup during neoadjuvant combination therapy. CD74 upregulation limited phagocytosis and stimulated M2 polarization. The CD74 blockade enhanced macrophage phagocytosis, resulting in decreased viability of cervical cancer cells in vitro and in vivo. This study unveils the dynamic microenvironment of UCC influenced by NACT and PD-1 blockade combination therapy, and targeting CD74 was a potential strategy to augment the therapy efficacy.

## Introduction

Cervical cancer continues to pose a significant burden on global health, necessitating the exploration of innovative treatment approaches to improve patient outcomes^1,2^. Despite advancements in conventional chemotherapy, there remains a need for more effective and targeted interventions^3^. Recently, immune checkpoint blockade has emerged as an encouraging strategy in cancer treatment, and ICIs have been approved for cervical cancer therapy. A large number of immunotherapies are being conducted in clinical trials with different potential targets, such as CTLA-4, 4-1BB, Tim-3, and PD-1/PD-L1^4,5^. Among these targets, the immunotherapy for programmed cell death protein 1 (PD-1) is more specific for cervical cancer. Pembrolizumab, nivolumab, balstilimab, cemiplimab, and cadonilimab have been used as second-line treatment for R/M CC patients^6^. However, more than half of the patients failed the treatment of PD-1 inhibitor^7^. Therefore, patients with advanced cervical cancer, especially those with recurrent and metastatic tumors, still need new therapies to improve outcomes.

Neoadjuvant chemotherapy (NACT) improved operability and subsequent treatment outcomes and may improve the long-term prognosis of resectable cervical cancer^8^. Platinum-based NACT showed a response rate of approximately 60% for patients with locally advanced cervical cancer^9^. Cisplatin interferes with DNA replication by cross-linking with DNA and is the most widely used chemotherapeutic agent in NACT for gynecologic malignancies. In the treatment process of many tumors, including gastric and ovarian cancer, general perspectives indicated that the combination of chemotherapy and immunotherapy can synergistically improve the effect of anti-cancer. This might be due to the reduced immune suppression of TME with the combination of chemotherapeutics and the enhanced elimination and recognition of tumors by the immune system^10,11^. PD-L1 blockade in combination with chemotherapy has already been applied in combination with chemotherapy in several clinical trials. In a phase 3 trial of patients with cervical cancer that recurred after first-line platinum-containing chemotherapy, survival was significantly longer with cemiplimab, a PD-1 antibody, than with single-agent chemotherapy^12^. The NCCN guidelines recommended pembrolizumab, a PD-1 monoclonal antibody, plus chemotherapy as the first-line therapy for PD-L1-positive advanced cervical cancer.

While the combination of neoadjuvant chemotherapy and PD-1 antibodies holds great promise, the underlying mechanisms and specific molecular players remain largely unidentified. One such molecule that has gained attention in the context of immune regulation of cancer is CD74. CD74 is a non-polymorphic type II transmembrane glycoprotein that has been shown to be responsible for antigen presentation, endocytic maturation, and cell migration ^13,14^. Furthermore, it is also considered a receptor for macrophage migration inhibitory factor (MIF), resulting in changes in macrophage function and polarization^15^. Studies have shown that CD74 activation could result in an M2 shift of macrophages^16^. Recently, CD74 has gradually attracted attention as a prognostic factor and therapeutic target for malignant tumor patients ^1,17–19^. However, its precise involvement in cervical cancer progression remains underexplored.

In this study, we performed single-cell RNA sequencing (scRNA-seq) analysis with the biopsy specimen from patients before chemotherapy, after NACT, and after receiving NACT and PD-1 blockade combination therapy, respectively. We revealed evidence of a transcriptomic evolution within the epithelial cells across distinct stages of the treatment process. Furthermore, we compared the dynamic alteration in the tumor microenvironment and found that CD74 ^high^ macrophages communicate frequently with immunoreactive Epi 3 subgroups. CD74 inhibited macrophage phagocytosis and induced M2 polarization, unfavorable to NACT combination immunotherapy for cervical cancer. Anti-CD74 Ab combined therapy in vitro and in vivo showed an improved anti-tumor effect. Together, our study comprehensively dissects the tumor microenvironment of UCC with NACT and PD-1 blockade combination at single-cell resolution and proposes that targeting CD74 could further enhance NACT treatment efficacy.

## Results

### Cellular dynamics and tumor microenvironment heterogeneity in cervical cancer receiving NACT and PD-1 blockade therapy

We collected nine tissues from three cervical cancer patients by puncture biopsy or surgery. The patient T3 and T4 received NACT and PD-1 blockade therapy before surgical treatment. We collected samples before (T3, T4) and after (T3a, T4a) NACT treatment and samples received PD-1 antibody-combined treatment (T3b, T4b, T4c). Samples before (T2) and after (T2a) NACT treatment of the patient T2 are also collected (Fig.1A). We analyzed 46950 cells in total and normalized the transcriptome expression profile. After annotation, 24 clusters were identified first and combined into seven, including cancer cells, MSC, Fibroblast, Endothelial cells, T cells, macrophages, and B cells (Fig.1B and S1). The relative cell ratios of the seven cell populations and UMAP projection of cells in the nine samples are shown in Fig.1 (Fig.1C and D). The cancer cell proportion is decreased after NACT and PD-1 antibody treatment. The immune cell type proportions showed tumor microenvironment heterogeneity (Fig.1E and F). We summarized the cell proportion during different treatment stages and found that the B cell proportion was significantly increased. Marker genes of every cell type were clustered in the heatmap (Fig.1G). Top marker genes of epithelial cancer cells include EPCAM, CDH1, and KRT8^20^. The Mesenchymal Stem Cell (MSC) cluster was identified with HSPB6, SERPINI1, and THY1^21^. Marker genes of fibroblast (DCN^22^), endothelial cells (VWF^23^), T cells (CD3D^24^), macrophages (CD14^25^), and B cells (CD79A^26^) were specifically expressed in their cluster (Fig.1H-J).

**Figure 1.**
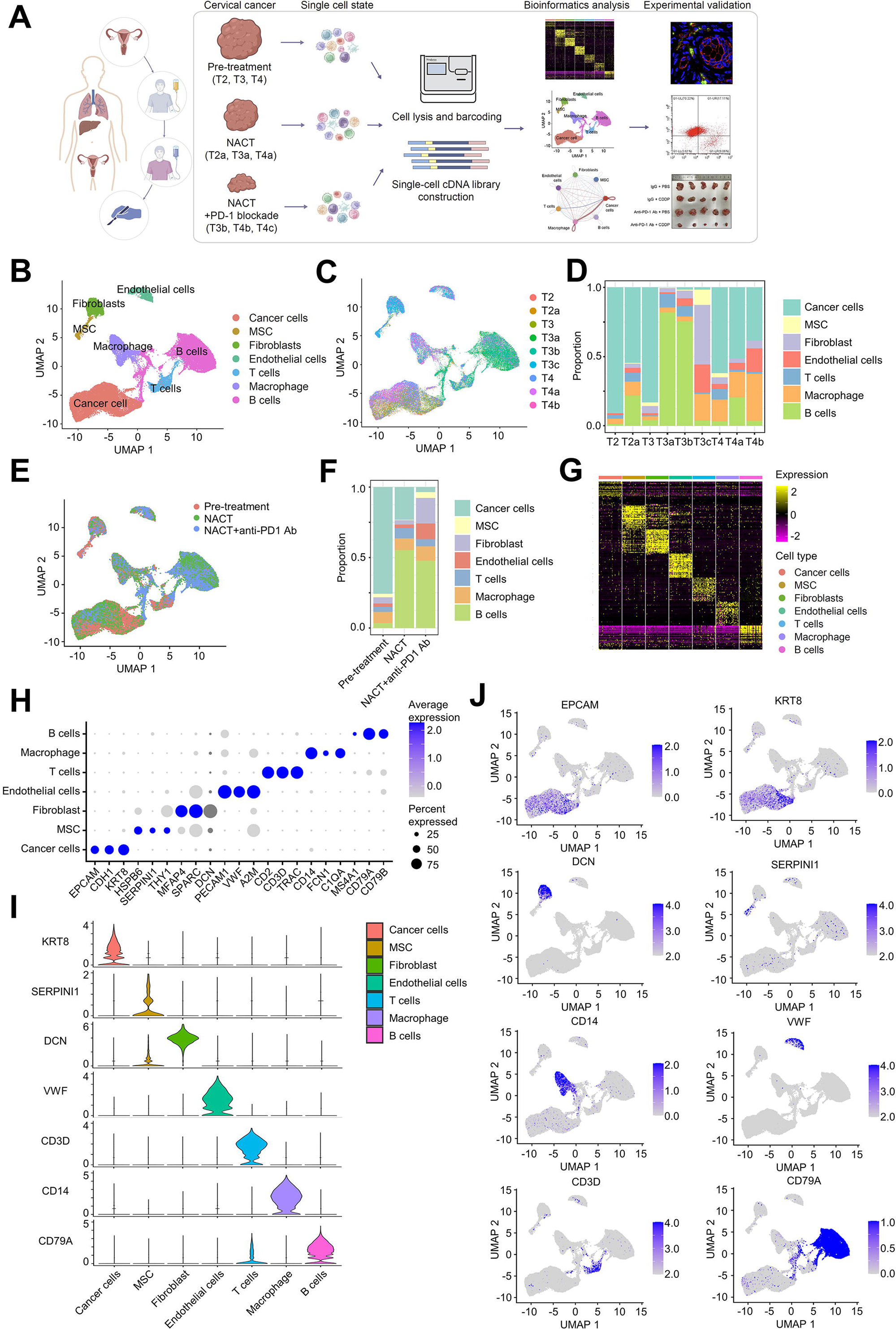
Study design and diverse cell types in cervical cancer receiving NACT and PD-1 antibody delineated by single-cell transcriptomic analysis. (A) The overall study design of the sample collection, scRNA-seq library construction, bioinformatics analysis, and experimental validation. The flowchart was created with biorender.com. (B) The Uniform Manifold Approximation and Projection (UMAP) plot demonstrates the cell types in cervical cancer tissues. (C) UMAP projection colored by each sample. (D) Proportions of assigned cell types in each sample are summarized in the stack bar plot. (E) UMAP projection colored by different stages of NACT combined anti-PD-1 Ab therapy. (F) The stack bar plot summarizes the cell type proportion of samples from different NACT combined anti-PD-1 Ab therapy stages. (G) Heatmap showing expression levels of marker genes in each cell type. (H) Dot plot of average expression and expression percentage of canonical marker genes for seven cell types. (I) Violin plots display representative marker expression across the cell types identified in cervical cancer. The y-axis shows the normalized read count. (J) UMAP plots of marker gene expression for cell type identification. The legend shows a color gradient of the normalized read count.

### Transcriptional evolution of epithelial cervical cancer cells during neoadjuvant combination therapy

As the dominant cell type, we supposed that epithelial cancer cells activated signature and pathway changes. The epithelial cells were subset from the mixed cells and reduced the dimension into six subclusters (Fig. 2A and S2A). Epi 1 proportion was decreased, and Epi 3 proportion was increased in post-platinum treatment groups, especially the T4 patient (Fig. 2B and S2B). Activated pathways in each cell subgroup were evaluated with the RRA algorithm. Epithelial subgroup 4 was enriched in the mitotic spindle, G2M checkpoint, E2F targets, and adipogenesis pathways. Epithelial subcluster 3 was immunoreactive, and IFN-gamma, IFN-alpha, and complement pathways were upregulated. Epithelial subgroup 2 was related to hypoxia, angiogenesis, apoptosis, and inflammatory response (Fig. 2C). Signal enrichment of mitotic spindle, IFN-alpha, glycolysis, and oxidative phosphorylation pathways, and others were consistent with UMAP distribution of corresponding subclusters (Fig. 2D and S2C). Pseudo-time analysis reveals the evolutionary process from Epi 4 to Epi 1 and Epi 2. The final destination is Epi 3 and Epi 5 (Fig. 2E). Timing of drug combination therapy correlates with Pseudotime. The pre-treatment group was the start, and the NACT and anti-PD-1 Ab group located dominant at the end (Fig. 2F). Monocle2 analysis was also used to rearrange the cells without UMAP characteristics and verify the consistency between the pseudo-time and timing of drug combination therapy (Fig. 2G-H and S2D). The cell population of pseudo-time state 2 was post-NACT treatment specific (Fig. S2E). Mitotic-related gene expression, including CDKN3, HMGB2, TOP2A, and CCNB1, was decreased along the pseudo-time. Immune response-related gene expression, including CD274, CD36, TNFSF10, CD69, TGFBI, and IFI27, was increased during the mid-term of the pseudo-time time series. In the end, ESR1, AGR3, APX8, VIM, MMP7, IGFBP8, XBP1, CD55 and MUC16 were upregulated (Fig. 2I and S2F). In all, NACT and anti-PD-1 Ab increase the immune reaction of the epithelial cells.

**Figure 2.**
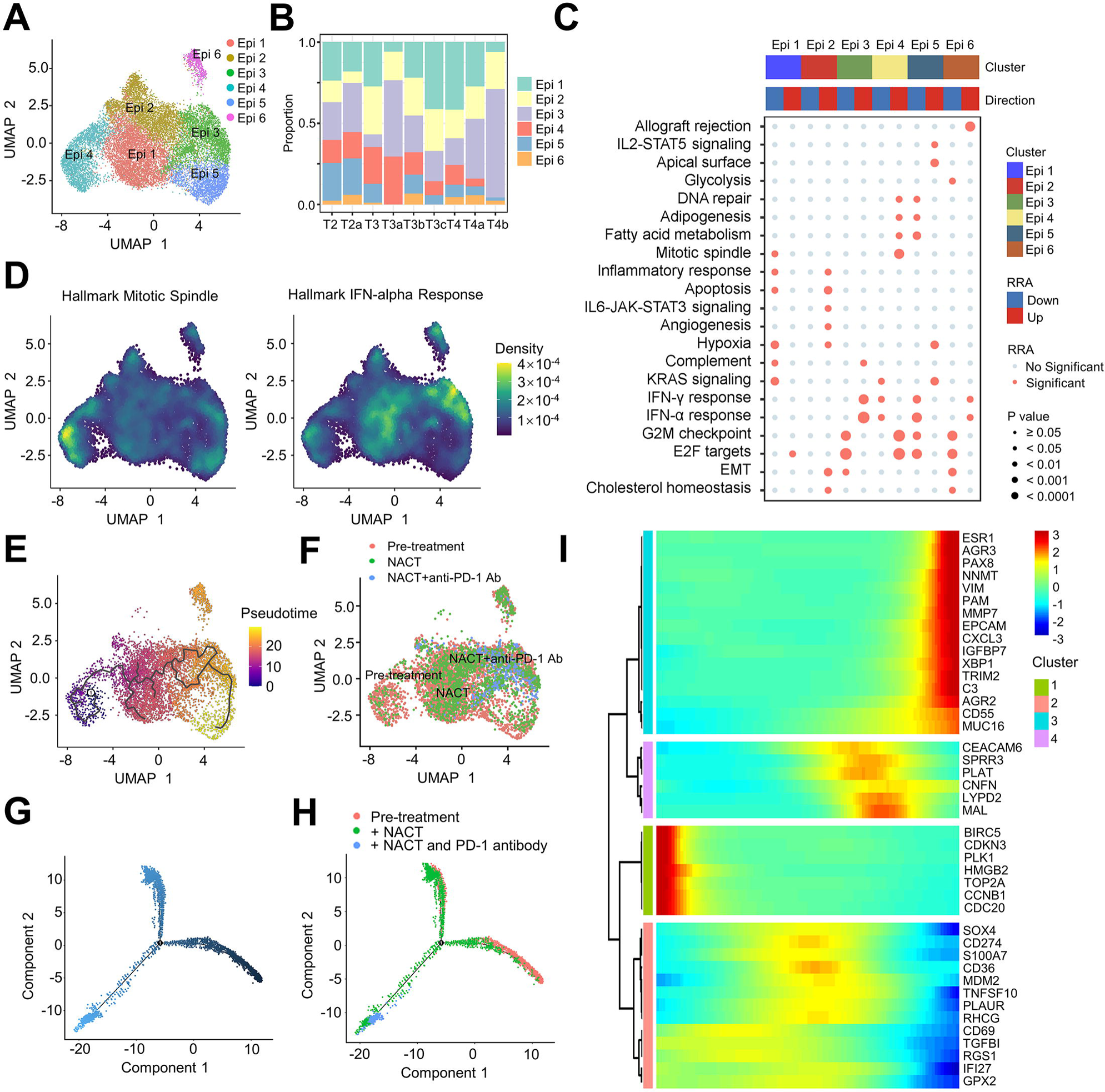
Cancer cell pseudo time analysis of cervical cancer receiving NACT and PD-1 antibody treatment. (A) UMAP projection of 6 subgroups generated from sub-clustering epithelial cancer cells. (B) Proportions of epithelial subgroups in each sample are summarized in the stack bar plot. (C) Pathway enrichment with RRA algorithm with top 100 signature genes specific to each epithelial subgroup. (D) UMAP projection of mitotic spindle and IFN-alpha response pathways density. (E) The unsupervised transcriptional trajectory of epithelial cells from Monocle (version 3), colored by pseudo-time. (F) UMAP projection of epithelial cell subgroups colored by different stages of NACT combined anti-PD-1 Ab therapy. (G) The unsupervised transcriptional trajectory of epithelial cells from Monocle (version 2), colored by pseudo-time. (H) Pseudo-time of epithelial cells inferred by Monocle (version 2) colored by different stages of NACT combined anti-PD-1 Ab therapy. (I) Heatmap showing the differential gene expression patterns in each state along the pseudo-time of NACT combined anti-PD-1 Ab therapy, clustered into four groups according to expression pattern.

### Epithelial cancer cell-immune cell communication changes dynamically during neoadjuvant combination therapy

To investigate the interaction between the cancer cells and the immune cells, we perform the cell communication analysis of the seven types of cell clusters. The total number and strength of inferred interactions increased after NACT treatment while decreasing after combined NACT and anti-PD-1 Ab treatment, indicating the potential immune activation function of NACT (Fig. 3A and 3B). We determined the number and strength of the interactions between each cell type. Communication between cancer cells, T cells, and macrophages was apparent (Fig. 3C and S3). Mainly, the cancer cell-T cell interaction strength decreased. In contrast, the interaction strength of cancer cell-macrophage increased after NACT treatment (Fig. 3D). The activated cancer cell-macrophage communication was silenced after the following period of NACT and PD-1 antibody treatment (Fig. 3E). Then, we determine the pathways and families underlying the global communication change. In terms of macrophage communication strength, we found that MHC-I and immune checkpoint signaling pathways significantly increased (Fig. 3F). The interaction strength of the immune checkpoint was diminished after combining with anti-PD-1 Ab (Fig. 3G). The PD-1 and PD-L1, immune checkpoint ligand-receptor interactions, participate in immune escape in cervical cancer. Though PD-1 failed to detect in scRNA-seq data, we investigated the ligand PD-L1 (CD274) expression in the cancer cell subcluster. CD274 was upregulated in immunoreactive Epi 2 and Epi 3 after NACT treatment (Fig. S3B-E). Pseudo-time analysis confirmed that CD274 is expressed at the mid-to-late temporal stage (Fig. S3F-G). The CDDP treatment did not increase PD-1 expression, while anti-PD-1 Ab decreased PD-1 expression in vitro and in vivo (Fig. S3H-J).

**Figure 3.**
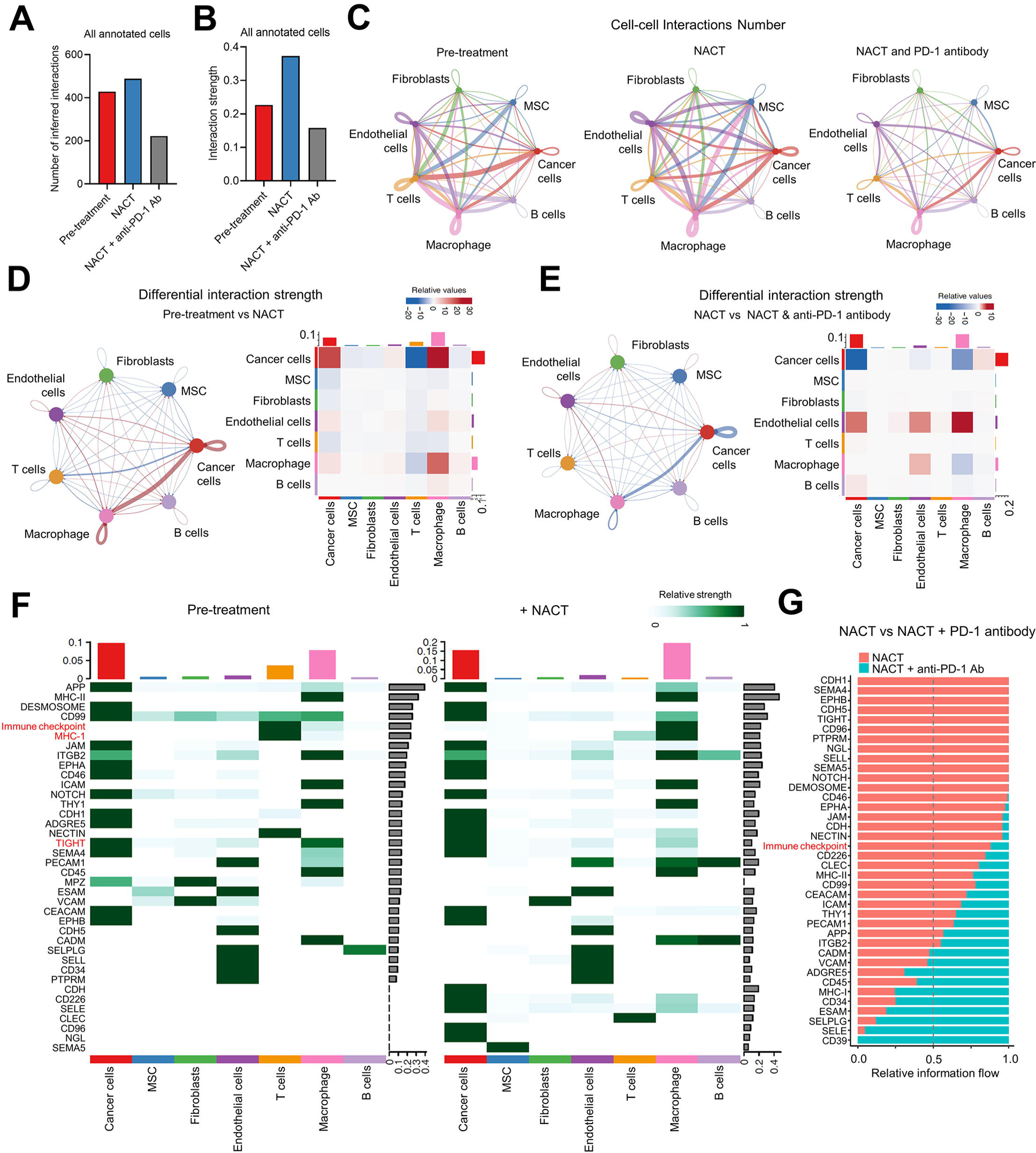
Cell communication analysis of tumor microenvironment after receiving NACT and PD-1 antibody treatment. (A, B) The number and strength of inferred interactions among seven types of cells before and after NACT treatment and the anti-PD-1 Ab combination. (C) The number of significant ligand-receptor pairs between any pair of two cell populations. The edge width is proportional to the indicated interaction number of ligand-receptor pairs. (D, E) The differential interaction strength between any pair of two cell populations before and after NACT treatment (D) and the anti-PD-1 Ab combination (E). The edge width and heatmap color are proportional to the normalized interaction strength. (F) The heatmap shows the relative interaction strength of 38 significant ligand-receptor signaling pathways belonging to each cell type. Left, the pre-treatment group. Right, the NACT treatment group. (G) Statistically significant signaling pathways were ranked based on their differences of overall information flow within the inferred networks between the NACT group and the anti-PD-1 Ab combination group.

### The therapeutic effect of anti-PD-1 Ab in combination with platinum drugs in cervical cancer partly depends on macrophages

We verified the combination effect of anti-PD-1 antibody and platinum drugs in vitro and clarified the necessity of macrophage-tumor cell communication for a combined therapeutic role. First, we treat the cervical cancer cell line SiHa and HeLa with IgG, CDDP, anti-PD-1 Ab, and combined CDDP and anti-PD-1 Ab. CDDP leads to increased apoptotic cell percentage (Fig. S4A-B). Besides, decreased cell viability and migration ability were detected with CCK8 and transwell assay after CDDP treatment. However, anti-PD-1 Ab did not directly affect cervical cancer cells, and combined treatment did not show any difference (Fig. S4C-E). When indirect coculture HeLa cervical cancer cells with macrophages, CDDP combined with anti-PD-1 Ab increased apoptotic cells compared with the CDDP group (Fig. 4A-B). The anti-PD-1 Ab combination also decreased the cell viability and migration ability (Fig. 4C-E). Then, we cocultured THP1-derived macrophages with cervical cancer cells SiHa. The results indicated CDDP inhibits macrophage phagocytosis ability after CDDP treatment (Fig. 4F). We also constructed a tumor subcutaneous tumorigenesis model with mice cervical cancer cell line TC-1 in immunocompetent C57 mice. The results showed that anti-PD-1 Ab elevated the inhibitory effect of CDDP on tumor growth (Fig. 4G-H).

**Figure 4.**
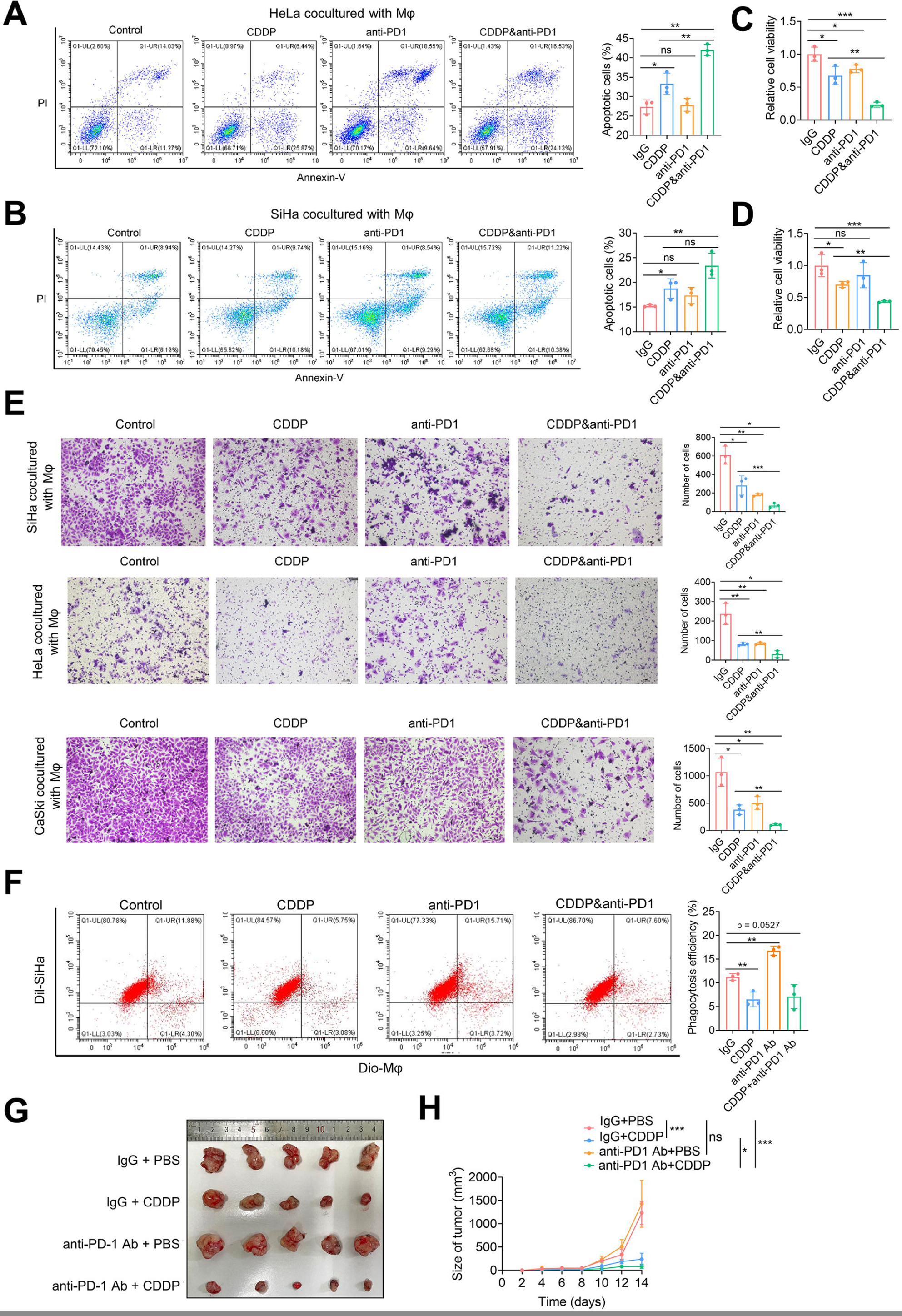
The therapeutic effect of PD-1 in combination with CDDP in cervical cancer is dependent on macrophages. (A, B) HeLa (A) or SiHa (B) cells cocultured with THP-1-derived macrophages were treated with anti-PD-1 Ab and/or CDDP. The upper right and lower right quadrants indicate the percentage of late or early apoptotic cells in the total measured cell population. Statistical graphs and representative experiments were shown. (C, D) The relative cell viability of HeLa (C) and SiHa (D) cells cocultured with THP-1-derived macrophages. (E) The migration ability of the cervical cancer cells cocultured with THP-1-derived macrophages was determined by transwell assay. The number of cells stained in purple indicates the level of migration ability. (F) Phagocytosis assay was performed using macrophages derived from THP-1 cocultured with SiHa. Phagocytosis efficiency was quantified by the percentage of Dil and Dio double fluorophore-positive THP-1-derived macrophages. (G) The representative image of the subcutaneous tumor. TC-1 cells were subcutaneously inoculated into C57BL/6J mice. Each group consisted of five mice and was treated with CDDP and/or anti-PD-1 Ab separately. (H) Growth curve of subcutaneous tumors in mice. The p-value was obtained by a two-tailed unpaired Student’s t-test, and the results are presented as the mean ±SD. *p<0.05, **p < 0.01, ***p<0.001

### Distinct subgroups of macrophages and their communication with epithelial cancer cell subgroups in cervical cancer

Macrophage-cancer cell communication changed along with the NACT and anti-PD-1 antibody treatment period. The macrophages were further divided into seven subgroups to determine how this alteration affects combination therapy (Fig. 5A and S5A). Each subgroup showed specific marker genes, and the Macro 3 and 5 subgroup proportion slightly increased after NACT treatment (Fig. 5B-C and S5B). The Macro 1 subgroup was enriched in complement-related immune response and neutrophil degranulation, and the Macro 4 subgroup was related to mitosis (Fig.S5C). Cell communication analysis was conducted between six epithelial subgroups and seven macrophage subgroups. The overall interaction strength increased after NACT treatment (Fig. 5D). Compared with the pretreatment group, the Epi 3 subgroup increased communication strength with the Macro 1-5 subgroup in the NACT group while decreased after anti-PD-1 treatment (Fig. 5E-G and S5D). The ligand-receptor families with increased interaction strength included immune checkpoint, CD226, CD39, and SN (Fig. 5H). NACT-induced differential interaction strength gradually decreased from Macro 1 to Macro 7. Thus, the seven subgroups were divided into three and two groups, respectively, to identify the decreasingly expressed marker genes (Fig. 5I). The Strong, Moderate, and Weak interaction strength group corresponds to UMAP clustering with clear demarcation (Fig. 5J). CD74, GRN, NPC2, CYBA, SLC40A1, and RAB42 were significantly in the Strong interaction strength group (Fig. 5K). Enriched pathways of the Strong interaction strength group and Macro 1 included phagocytosis, antigen processing and presentation, neutrophil degranulation, and PD-1 signaling (Fig. 5L and S5E-F). Collectively, the macrophage subgroup that interacted with cancer cells during NACT therapy showed a phagocytosis signature related to the immune checkpoint.

**Figure 5.**
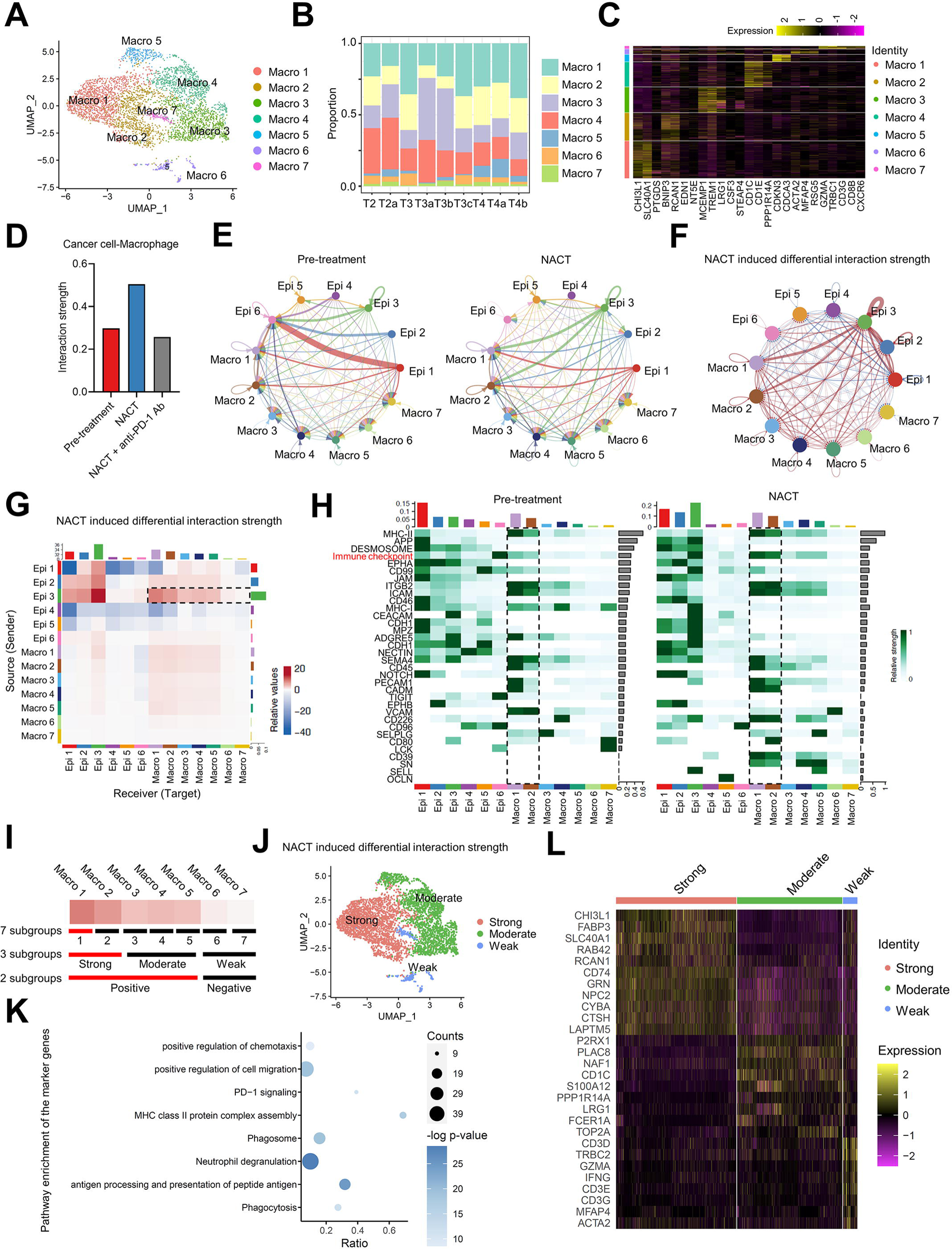
Macrophage subgroups and cell communication with epithelial subgroups. (A) UMAP projection of seven subgroups generated from sub-clustering macrophages. (B) Proportions of macrophage subgroups in each sample are summarized in the stack bar plot. (C) Heatmap showing expression levels of marker genes in each macrophage subgroup. (D) The strength of inferred interactions among epithelial cancer cell and macrophage subgroups before and after NACT treatment and the anti-PD-1 Ab combination. (E) The interaction strength of significant ligand-receptor pairs between any pair of two epithelial cancer cell and macrophage subgroups. The edge width is proportional to the indicated interaction strength of ligand-receptor pairs. (F-G) The circular network plot (F) and heatmap (G) of the differential interaction strength between any pair of two epithelial cancer cell and macrophage subgroups before and after NACT treatment. (H) The relative interaction strength heatmap of 34 significant ligand-receptor signaling pathways belonging to each epithelial cancer cell and macrophage subgroup. Left, the pre-treatment group. Right, the NACT treatment group. (I) Grouping method design for macrophage subgroups to screen for differential marker genes. Seven macrophage subgroups were further divided into Strong (Macro 1 and 2), Moderate (Macro 3, 4, and 5), and Weak groups (Macro 6 and 7). The Positive (Macro 1-5) and Negative (Macro 6-7) groups were divided according to the differential interaction strength after NACT treatment. Marker genes of the subgroups colored in red were identified. (J) UMAP projection of the Strong, Moderate, and Weak subgroups. Colored by different subgroups of differential interaction strength. (K) Pathway enrichment of the genes shared by the Strong and Macro 1 subgroups. (L) Heatmap showing expression levels of marker genes in the Strong, Moderate, and Weak subgroups.

### CD74 expression in macrophages is elevated in subgroups with strong communication strength and after combination therapy

To identify the dominant genes in the variation of cancer cell-macrophage communication strength, we took intersections in the Macro 1, Strong, and Positive subgroups. We obtained 18 candidate genes (Fig. 6A). The candidates, including CD74, GRN, CYBA, CTSH, and others, showed the highest expression in the Strong subgroup and lowest expression in the Weak subgroup (Fig. 6B and S6A). We compared the expression specificity of the 18 candidates in different subgroups and samples and found that post-NACT upregulation of CD74 was consistent across all patients (Fig. 6C and S6B). CD74 expression increased along the pseudo-time of the macrophage subgroup and was highest in the Strong subgroup (Fig. 6D-G and S6C-E). Then, we verified CD74 expression in the patient tissues. CD74+ ratio increased in CD68+ macrophages, and NACT-induced CD74 upregulated in T2, T3, and T4 patients (Fig. 6H and S6F). In vitro, macrophages cocultured with CaSki and HeLa cells showed increased CD74 mRNA and protein levels after CDDP treatment. Though anti-PD-1 Ab did not result in CD74 upregulation, the RNA and protein expression level in the combination therapy group was about two times higher than that of CDDP treatment alone (Fig. 6I-J and S6G-H). In the subcutaneous graft tumor model, immunofluorescence assay indicated that the CD74+/CD68+ cells in NACT treatment groups were more than in control and anti-PD-1 Ab groups (Fig. 6K). The solid tumor was then isolated into single cells, and the flow cytometry results showed that the CD74+ cells ratio of macrophages increased in the NACT group (Fig. 6L and S6I). In all, CD74 upregulation is an important effect induced by NACT combined with immune therapy and participated in macrophage-cancer cell communication.

**Figure 6.**
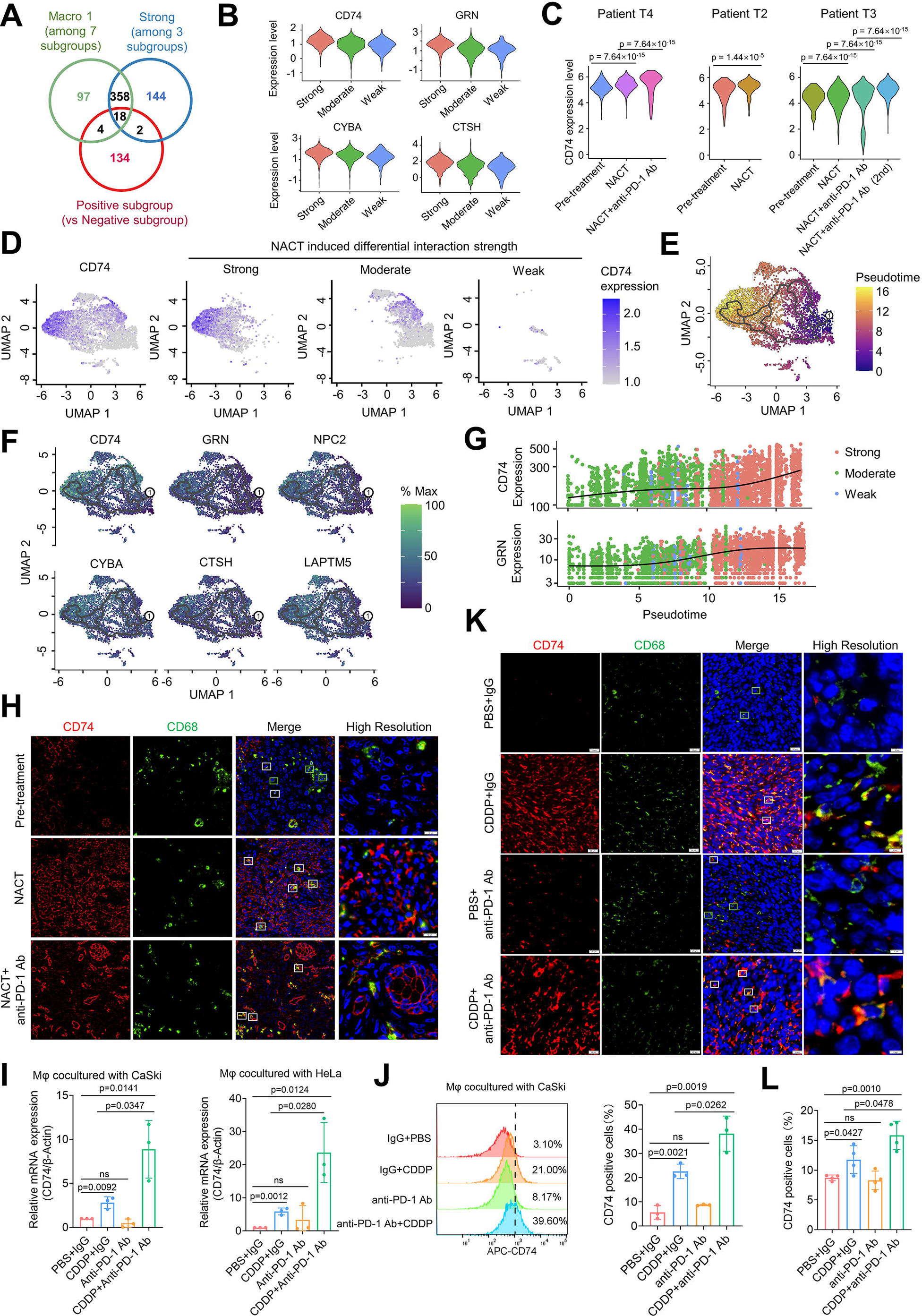
CD74 expression level positively correlated with the strength of macrophage-cancer cell communication and increased after NACT treatment. (A) Venn diagram of the 18 candidate genes shared by the Macro 1, Strong, and Positive subgroups. (B) Violin plot of the CD74, GRN, CYBA, and CTSH expression in the Strong, Moderate, and Weak subgroups. (C) The normalized CD74 expression level after pre-treatment, NACT, and anti-PD-1 Ab combination therapy (NACT+anti-PD-1 Ab) in T2, T3, and T4 patients. The patient T3 samples received two rounds of NACT and anti-PD-1 Ab combination therapy were analyzed. (D) UMAP projection of the CD74 expression in macrophage subgroups. The expression plot was split based on the Strong, Moderate, and Weak subgroups. (E) The unsupervised transcriptional trajectory of macrophages from Monocle (version 3), colored by the pseudo-time. The circle labeled 1 indicated the start of the trajectory. (F) UMAP projection of the normalized expression of CD74, GRN, NPC2, CYBA, CTSH, and LAPTM5 in macrophage subgroups. The gray curve indicates the pseudo-time trajectory. (G) The CD74 and GRN expression dynamic change along with the pseudo-time, colored by the Strong, Moderate, and Weak subgroups. (H) The immunofluorescence assay showed the CD74 (red) and CD68 (green) expression in the tissues of patient T4 receiving NACT and anti-PD-1 Ab combination therapy. The green boxes represent CD74 negative macrophages, and the white boxes represent macrophages expressing CD74. (I) The CD74 mRNA level in HeLa or CaSki cells cocultured with macrophages derived from THP-1 cells. Anti-PD-1 Ab and CDDP were used to establish a model of NACT combined with immunotherapy in vitro. (J) The percentage of CD74-positive macrophages cocultured with CaSki cells was detected by flow cytometry. (K) The immunofluorescence assay showed the CD74 (red) and CD68 (green) expression of subcutaneous tumors treated with CDDP and/or anti-PD-1 Ab. (L) Flow cytometry revealed the number of CD74-positive macrophages in single-cell suspensions isolated from mice subcutaneous tumors. The p-value was obtained by a two-tailed unpaired Student’s t-test, and the results are presented as the mean ±SD.

### CD74 inhibits phagocytosis and induces M2 polarization of macrophage

To explore the effect of NACT-induced CD74 upregulation, we use siRNA targeting CD74 and CD74 overexpress plasmid transfected into THP-1-induced macrophages. Phagocytotic efficiency increased after CD74 knockdown and decreased when CD74 was overexpressed (Fig. 7A-B and S7A-D). We also blocked the CD74 with anti-CD74 Ab and found increased phagocytotic efficiency (Fig. 7C). The CD74 mRNA level was unaffected after anti-CD74 Ab treatment, while protein expression was slightly decreased (Fig. S7E-F). We speculated whether CD74-induced phagocytosis change was related to macrophage M1/M2 polarization. M1/M2 ssGSEA score of the Strong, Moderate, and Weak subgroups was analyzed with scRNA-seq data of the macrophage population. M2 ssGSEA score was highest in the Strong subgroup (Fig. 7D). M1/M2 ssGSEA score decreased after each period of NACT combination therapy, indicating macrophage transformation from M1 to M2 (Fig. 7E). CD74 signal intensity distribution was similar to that of M2 score and was opposite to that of M1 score (Fig. 7F-G). Flow cytometry results indicated that CD74 overexpression led to CD86 downregulation and CD206 upregulation. Knockdown of CD74 had the opposite impact (Fig. 7H-K). Knockdown CD74 reversed NACT-induced loss of phagocytosis ability in vitro (Fig. 7L and S7G-H). Thus, we attempt to apply anti-CD74 Ab to improve the combination therapeutic effect. We use cervical cancer SiHa cells cocultured with macrophages and anti-CD74 Ab restored phagocytic impairment due to NACT (Fig. 7M). In vivo, we use CD74 humanized BALB/c-hCD74 mice to determine the anti-tumor ability of anti-CD74 Ab combined with NACT treatment (Fig. 7N-O). Spleen-derived macrophages were isolated from humanized mice and co-cultured with mice cervical cancer TC-1 cells. The phagocytotic efficiency of the anti-CD74 Ab combined treatment group was higher than that of traditional NACT treatment alone (Fig. 7P). Moreover, MIF, the ligand for CD74, is not explicitly expressed across any cancer cell subpopulations and samples (Fig. S7I-L). Besides, CD74 was irrelevant to the PD-1 expression of macrophages (Fig. S7M-N). In summary, the anti-CD74 Ab combined NACT therapy improves the anti-tumor effect and provides a promising targeting strategy for cervical cancer treatment.

**Figure 7.**
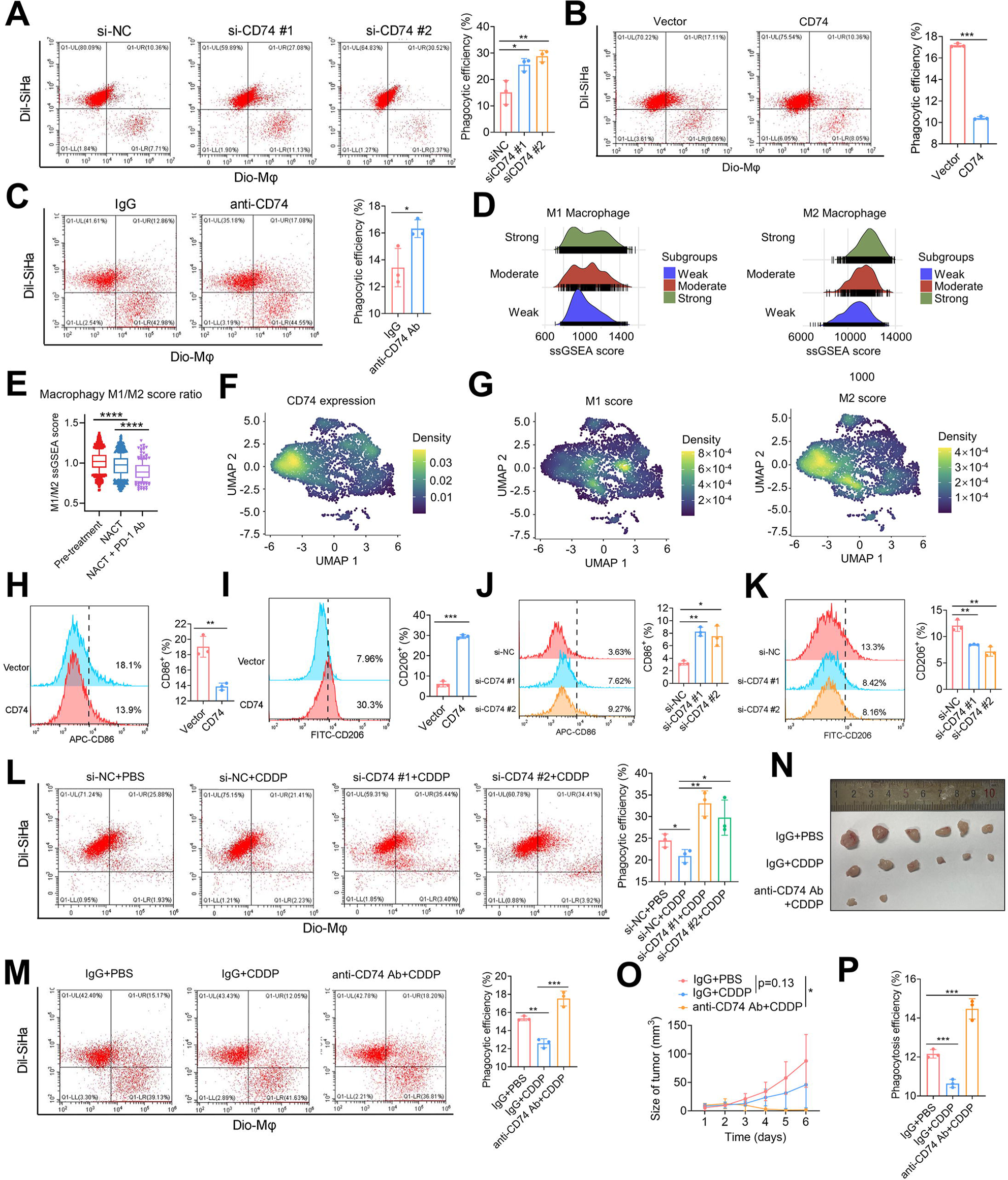
CD74 inhibits phagocytosis and promotes M2 polarization of macrophage induced by NACT. (A, B) Phagocytosis ability detected by flow cytometry. Macrophages derived from THP-1 cells were co-cultured with SiHa cells knockdown (A) or overexpressing (B) CD74. Phagocytosis efficiency was quantified by the percentage of double fluorophore-positive macrophages in the upper right quadrant. (C) The phagocytic function of macrophages co-cultured with SiHa cells treated with anti-CD74 was measured by flow cytometry. (D) Density plot of the M1 and M2 ssGSEA score in the Strong, Moderate, and Weak subgroups. (E) M1/M2 ssGSEA ratio of the macrophages in the pretreatment, NACT, and anti-PD-1 Ab combination group. (F, G) UMAP projection of the normalized density of CD74 expression (F), M1 score, and M2 score (G) in the macrophage subgroups. (H, I) CD86 (H) and CD206 (I) expression of THP-1-derived macrophages after CD74 overexpression. CD86 is the surface marker of M1 macrophages, while CD206 is the surface marker of M2 macrophages. (J, K) CD86 (J) and CD206 (K) expression of THP-1-derived macrophages after CD74 knockdown. (L) Phagocytosis efficiency of THP-1-derived macrophages cocultured with SiHa cells. CDDP-induced phagocytic function loss of macrophages was rescued by knocking down CD74. (M) Flow cytometry detected the differences in phagocytic function among the control, the CDDP, and the anti-CD74 Ab combined with the CDDP group. THP-1-derived macrophages were cocultured with SiHa cells. (N) The representative image of the subcutaneous tumor model. Mouse cervical cancer cells TC-1 were subcutaneously inoculated into immunocompetent CD74 humanized BALB/c-hCD74 mice. Each group consisted of 6 mice receiving CDDP or CDDP combined with intraperitoneal injected anti-CD74 Ab. Tumors of four mice in the drug-combination group became nonpalpable on Day 4 after administration. (O) Growth curve of subcutaneous tumors in mice. (P) Phagocytosis assay was performed using macrophages isolated from the mice’s spleen and mouse cervical cancer cell line TC-1. Phagocytosis efficiency was quantified by the percentage of Dil and Dio double fluorophore-positive THP-1-derived macrophages. The p-value was obtained by a two-tailed unpaired Student’s t-test, and the results are presented as the mean ±SD. *p<0.05, **p < 0.01, ***p<0.001

## Discussion

Platinum-based NACT was widely used in cervical cancer pre-surgical treatment. Immune therapy combined with NACT in cervical cancer treatment is still in clinical trials (NCT04799639), and there was a lack of molecular biological evidence. Here, we collected sequential samples from cervical cancer patients receiving NACT combined with anti-PD-1 Ab treatment before surgery. Single-cell RNA-seq revealed complicated proportion and communication alteration in cancer cells, fibroblasts, and other immune cells. We comprehensively investigated the dynamic expression profiles of macrophage and epithelial cell subpopulations during various therapy stages. Our data provides a valuable source for clarifying the underlying mechanism and regulatory network of combination therapy, enabling more discovery of new targets to improve therapeutic efficacy and contributing to patient staging and precise treatment.

Previous scRNA-seq studies focus on the heterogeneity of human cervical squamous cell carcinoma initiation and progression and capture the overall picture ^27,28^. Specific subpopulations of cervical cancer, such as the HPV-reprogrammed keratinocyte subpopulation, were identified to provide evidence for tumor virus infection and cancer evolution^29^. The radiochemotherapy-induced innate immune activation and MHC-II upregulation in cervical cancer was also revealed by scRNA-seq^30^. Single-cell dissection of T cell subpopulation in cervical cancer tissues showed that targeting IDO1 and immune checkpoint inhibitors could improve the therapeutic efficacy^31^. Our study focused on macrophage-cancer cell communication during the NACT in combination with anti-PD-1 Ab therapy and identified CD74 as an unfavorable factor. Besides, we found candidate markers of epithelial cancer cell population upregulated in the later stages of treatment, including ESR1, AGR3, PAX8, NNMT, VIM, PAM, MMP7, CXCL3, IGFBP7, XBP1, AGR2, CD55, MUC16. Several markers inside the cancer cell population aroused by the immune response, including CEACAM6, SPRR3, CNFN, SOX4, S100A7, CD36, MDM2, TNFSF10, PLAUR, CD69, TGFBI, IFI27, GPX2, warranting further investigation for their roles in combination therapy.

Our study elucidates the immune microenvironmental basis for PD-1 to enhance the efficacy of NACT. After the first round of NACT, there is a significant decrease in cervical cancer cells and an increase in the percentage of immune cells, such as B cells, representing a higher degree of immune infiltration. Tumor cells respond to the immune system attack and activate immune response-related pathways. One of the crucial actions is to elevate the expression of CD274 and send a “don’t eat me” signal to immune cells. We found that the increased immune checkpoint signal mainly existed between macrophages and cancer cells. The macrophage subgroup receiving the strongest signal was related to the positive regulation of chemotaxis and antigen processing and presentation of peptide antigen, which may facilitate immune recognition and activation. Moreover, the macrophage subgroup showed an M2 polarization signature, which was reported to sensitize anti-PD-1 treatment^32,33^. The above results support the evidence that the anti-PD-1 Ab immune therapy combination leads to better outcomes of NACT in cervical cancer treatment.

However, we found that platinum-based NACT-induced changes were not all favorable to cervical cancer treatment. T cell-cancer cell communication decreased, and the proportion of T cells was not significantly changed, consistent with a previous study ^34^. Besides, though macrophage-cancer cell communication increases, the upregulated strength ligand-receptor interactions include immune checkpoint, MHC-1 family, and TIGHT. These pathways, except PD-1, will keep releasing defense signals to the immune system. In terms of macrophages themselves, the phagocytosis ability decreases after platinum-based NACT treatment. These results prevent further enhancement of the efficacy of combination therapy.

As a critical component in the tumor microenvironment, TAM recruitment is associated with malignant progression, therapy resistance, and ferroptosis in cervical cancer ^35,36^. However, the heterogeneity and M1/M2 polarization of TAMs have challenged the investigation of their communication with cervical cancer cells. In this study, we found the progressive upregulation of CD74 in macrophages and partly explained the diminished macrophages after neoadjuvant therapy. The expression level of CD74 correlates with communication strength through cell communication analysis. Previous studies found that the CD74/MIF signal pathway participates in cancer under pathological conditions, inflammation, and autoimmune diseases^37,38^. CD74 signaling is dominant in tumor-infiltrating macrophages in low-grade glioma, impedes microglial M1 polarization, and facilitates brain tumorigenesis^16,39^. We validated that CD74 contributes to macrophage M2 polarization and weakened phagocytosis ability in cervical cancer, which plays an obstructive role in immunotherapy resistance ^40^. Our research uses siCD74 and anti-CD74 Ab to counteract M2 polarization. Anti-CD74 combination therapy reverses macrophages from M2 to M1 morphology, increasing phagocytosis anti-tumor activity^41^. M1 polarized macrophages may increase anti-tumor immune responses’ sensitivity and response rate through cross-talk with other immune cells. Targeting CD74 was a potential strategy to further improve NACT and PD-1 blockade combination therapy.

Though the study found potential targets for improving neoadjuvant therapy efficacy, the current study has limitations and requires further research. First, immune therapy combined with NACT for cervical cancer treatment is still under clinical trial, and the sequential samples of the same patient before and after neoadjuvant therapy are precious. So, the sample size in this study was small, and a large sample size would help to validate our findings further. Second, our study focused on the macrophages, while the tumor microenvironment was complex and dynamic. Subcluster of T and B cells and their communication with cancer cells need further investigation. Third, the molecular profile of the epithelial cancer cells during the NACT and anti-PD-1 Ab treatment needs additional validation.

Our study provides a comprehensive interaction network of the tumor microenvironment, especially between macrophages and epithelial cancer cells, during the NACT and anti-PD-1 Ab treatment. We validated the efficacy of the combination therapy in vivo and in vitro and revealed that anti-PD-1 Ab improves the anti-tumor effect of NACT partly depending on macrophages. Moreover, CD74 upregulation in macrophages is a molecular signature in combination therapy. Targeting CD74 effectively enhances the anti-tumor effect of neoadjuvant therapy for cervical cancer.

## Methods

### Patients and sample collection

The clinical trial and experimental research were approved by the Research Ethics Committee of Shandong University Qilu Hospital (KYLL-2020B-029, KYLL-202111-153). 9 specimens were collected from stage IB3 and IIA2 (FIGO 2018^42^) female cervical cancer patients who were treated at the Department of Gynecology and Obstetrics, Qilu Hospital, Shandong University. The patients had not undergone prior treatment and received platinum-based NACT combined with PD-1 blockade therapy before radical hysterectomy. The pre-treatment (T2, T3, T4) and post-NACT (T2a, T3a, T4a) specimens were collected by puncture biopsy. The post-NACT combined with PD-1 blockade specimens (T3b, T4b, T4c) were collected at the time of surgical resection under the supervision of a qualified pathologist. All patients provided written informed consent.

### Tissue dissociation and preparation

Within 30 minutes after surgery, the newly collected tissues were preserved on ice in sCelLiveTM Tissue Preservation Solution (Singleron). The specimens were washed using Hanks Balanced Salt Solution (HBSS) and finely fragmented into small pieces. Singleron PythoN™ Tissue Dissociation System performed tissue digestion with sCelLiveTM Tissue Dissociation Solution (Singleron). The cell suspension was filtered through a 40 μm sterile strainer. 2 volumes of the GEXSCOPE® red blood cell lysis buffer (Singleron) were mixed and incubated at room temperature for 5-8 min. The mixture was then centrifuged and suspended in PBS (HyClone) for downstream library construction.

### Single-cell library construction

Single-cell library construction was performed by Singleron Matrix® Single Cell Processing System according to the instructions of the GEXSCOPE® Single Cell RNA Library Kits^43^. Single-cell suspensions with a concentration of 2×10^5^ cells/mL were loaded onto a microwell chip. Barcoding Beads from the microwell chip were collected and used for reverse transcription and PCR amplification. The cDNA was fragmented and ligated with sequencing adapters. Libraries were sequenced on the Illumina NovaSeq 6000 platform (PE150).

### Primary analysis of the sequencing data

Gene expression profiles were generated from the raw reads utilizing CeleScope (v1.5.2, Singleron) with default parameters. The process involved extracting barcodes and UMIs from R1 reads, followed by their correction. Adapter sequences and poly-A tails were trimmed from R2 reads, and the clean R2 reads were aligned to the GRCh38 (hg38) transcriptome using STAR (v2.6.1a)^44^. Uniquely mapped reads were assigned to genes using FeatureCounts (v2.0.1)^45^. Reads with the same cell barcode, UMI, and gene were grouped to generate the gene expression matrix for subsequent analysis.

### Quality control, dimension-reduction, and clustering

Seurat (4.3.0)^46^ was used for quality control, dimensionality reduction, and clustering under R (4.3.1). For each sample dataset, we filtered the expression matrix by the following criteria: 1) cells with a gene count less than 200 or with a top 2.5 % gene count were excluded; 2) cells with a top 2.5 % UMI count were excluded; 3) cells with over 20 % of UMIs derived from the mitochondrial genome were excluded; 4) genes expressed in less than 5 cells were excluded. After filtering, 46950 cells were retained for the downstream analyses. SCTransfrom function in the Seurat R package was used to minimize the batch effects and normalization. 3,000 highly variable genes were identified in each sample based on a variance-stabilizing transformation to generate an integrated expression matrix. Principle Component Analysis (PCA) was performed on the scaled variable gene matrix, and the top 15 principal components were used for clustering and dimensional reduction. Cells were separated into 24 clusters using the Louvain algorithm, setting the resolution parameter at 0.8. Cell clusters were visualized by using Uniform Manifold Approximation and Projection (UMAP).

### Cell type annotation and subtyping of epithelium and macrophages

The differential expressed genes of each cell subcluster were identified by the FindAllMarker function of Seurat (v 4.3.0)^47^ with the parameters logfc.threshold = 0.25, test.use = “wilcox”, min.pct = 0.1, only.pos = T. Cell types were annotated by SingleR with manual correction and identification. Canonical cell type markers for single-cell seq data were achieved from CellMakerDB, PanglaoDB, and recently published literature. To obtain a high-resolution map of Epithelial cancer cells and macrophages, cells from the specific cluster were extracted and reclustered for more detailed analysis following the same procedures described above and by setting the clustering resolution as 0.3.

### Cell-cell interaction analysis

Cell-cell interaction analysis was performed with Cellchat (v1.5.0)^48^. We added 42 extra receptor-ligand pairs of immune checkpoint pathways into the CellChatDB based on the previous study ^49^. The gene expression matrix was projected to proteome-proteome interaction networks to reduce the dropout effect of signaling genes. Due to the completeness of the series of samples and the stability of the data quality, the T4 sample was selected for preliminary analysis and other samples for validation. The number and strength of the inferred interactions were analyzed between seven main cell types. The differential interaction strength was compared between the Pre-treatment and NACT groups and between the NACT and the NACT & anti-PD-1 antibody groups. The relative interaction strength and information flow of the detailed receptor-ligand signal pathways were analyzed. Cell communication was also analyzed between six epithelial subgroups and seven macrophage subgroups. The epithelial and macrophage subgroups were merged to create a Cellchat object and analyzed following the same process.

### Pseudotime Trajectory Analysis

The differentiation trajectory of epithelial cancer cells and macrophage subtypes was reconstructed with the Monocle2 (v 2.24.1) and Monocle3 (v 1.2.9)^50,51^. For constructing the trajectory, highly variable genes (q-value < 0.01) were selected by the differentialGeneTest function, and dimension-reduction was performed by DDRTree. The trajectory visualization was conducted by the plot_cell_trajectory function in Monocle2. Monocle3 was also used in pseudotime trajectory analysis. UMAP projection was imported from Seurat Object. The trajectory was created with the learn_graph function. The subpopulation of cells in which the Pre-treatment group predominates was chosen as the starting point for the pseudotime trajectory. The trajectory corresponding to UMAP projections was visualized by the plot_cells function in Monocle3.

### Functional annotation and enrichment analysis

Pathway enrichment in each epithelial cancer cell subgroup was performed with the irGSEA package (v 1.1.3) (https://github.com/chuiqin/irGSEA). The pre-defined sets of genes in the MSigDB database were used for analysis. The rank aggregation algorithm (RRA) integrated the differential pathways from AUCell^52^, UCell^53^, singscore^54^, and ssGSEA^55^ and was used for visualization^56^. Pathway enrichment in each macrophage subgroup was performed with the clusterProfiler package (v 4.4.4)^57^. The top 100 marker genes in each subgroup were identified with the COSG package (v 0.9.0)^58^. Significant Reactome pathways (p-value < 0.05) were visualized with the dotplot function. Gene signatures of M1 and M2 macrophages used for ssGSEA enrichment were achieved from the previous study^59^. UMAP projection of pathway enrichment and relative expression was visualized by Nebulosa^60^.

### In vivo studies

C57bl-6j and BALB/cJGpt-Cd74em1Cin(hCD74)/Gpt mice were purchased from GemPharmatech (Nanjing, China). Animal studies were performed according to institutional guidelines, and all experiments in this study were approved by the Ethics Committee of the Qilu Hospital of Shandong University (Grant NO. DWLL-2023-030). A total of 2 × 10^5^ viable cells were injected into the right flanks of mice. Anti-mouse PD-1 (CD279)-InVivo (10 mg/kg, every other day, Selleck, China) and Cisplatin (7 mg/kg, every four days, MedChemExpress, China) were injected into the abdominal cavity of mice to build a NACT combined with PD-1 therapy model. Milatuzumab (anti-CD74) (15 mg/kg, every other day, MedChemExpress, China) was used to block the expression of CD74. Mice in the control group were intraperitoneally injected with InVivoMAb polyclonal Armenian hamster IgG (10 mg/kg, every other day, Bioxcell, USA) and PBS. Then, after 12 days, they were sacrificed, the tumors were dissected, and tumor weights were measured. Tumor sizes were measured using a Vernier caliper, and the tumor volume was calculated using the following formula: volume = 1/2 × length × width^2^.

### Immunofluorescence Staining

Multiple immunofluorescence kits (Immunoway, China) are used for multicolor immunofluorescence staining. For CD74 and CD68 staining, histological section and cells were stained with anti-CD74 antibody (Abcam, Cat#ab108393, 1:500; Abcam, Cat#ab289885, 1:500) and anti-CD68 antibody (CST, Cat#97778, 1:200). Then, the samples were treated with an HRP polypolymer secondary antibody (Anti-Rabbit/Mouse) and fluorescent chromogenic solution according to the kit instructions. A fluorescence microscope and confocal microscope were used to observe experimental results.

### Cell culture and establishment of a co-culture system

CaSki and THP-1 cells were cultured in 1640 Medium (GIBICO, US), and HeLa and SiHa cells were cultured in DMEM Medium (GIBICO, US). The medium contains 10 % fetal bovine serum (GIBCO, US) and 1% Penicillin-Streptomycin (100×) (Solarbio, US), and cells were cultured at 37 °C with a 5 % CO2 atmosphere.

THP-1-derived macrophages were obtained after phorbol ester treatment (PMA, Sigma, USA) (50 ng/ml) for 48h. CaSki, HeLa, and SiHa cells were cocultured with THP-1-derived M2 macrophages using Transwell insert (0.4 μm; Corning, USA). Each lower well of plates was plated with approximately 10,000 cervical cancer cells, and the upper well was plated with 5,000 THP-1-derived macrophages. TC-1 cells are mouse lung epithelial cells stably transfected with E6/E7. TC-1 cells are commonly used to construct animal models of cervical cancer. TC-1 was cultivated in DMEM medium (Gibco, USA) containing 10% fetal bovine serum (GIBCO, US) and 1% Penicillin-Streptomycin (100×) (Solarbio, US) at 37°C in a humidified 5% CO2 incubator. Cisplatin (30 μM, 4h, MedChemExpress, China) and Pembrolizumab (anti-PD-1 Ab) (20μg/ml, 24h, MedChemExpress, China) were used to act on a co-culture system to build a NACT combined with PD-1 therapy model.

### Flow cytometry

Macrophages derived from THP-1 cells and single-cell suspension from mouse tumors were collected and evaluated by flow cytometry. Single-cell suspension from mouse tumors or spleens were stained with Alexa Fluor 700 anti-mouse CD45 Antibody (Biolegend, USA, Cat#147716), APC anti-mouse F4/80 Antibody (Biolegend, USA, Cat#123115), PE/Cyanine7 anti-mouse/human CD11b Antibody (Biolegend, USA, Cat#101215), Alexa Fluor 488 anti-mouse CD74 (CLIP) Antibody (Biolegend, USA, Cat#151005) and PE anti-mouse CD279 (PD-1) Antibody (Biolegend, USA, Cat#135206). CD45 antibody is used to screen monocytes in cell suspension. In the CD45-positive cell population, the macrophages expressing both F4/80 and CD11b were gated to detect PD-1 and CD74 expression.

THP-1-derived macrophages were stained with the following fluorochrome-labeled antibody. FITC anti-human CD206 (MMR) Antibody (Biolegnd, USA, Cat#321104), PerCP/Cyanine5.5 anti-human CD163 Antibody (Biolegend, USA, Cat#326512), APC anti-human CD74 Antibody (Biolegend, USA, Cat#326811) and APC anti-human CD279 (PD-1) Antibody (Biolegend, USA, Cat#329908).

The apoptotic cells were detected using the Annexin V-FITC Apoptosis Detection Kit (Beyotime, China). Cells were harvested and resuspended in the Annexin V-FITC and PI binding buffer. The total percentage of each group containing early and late apoptosis was compared.

The stained samples were detected using CytoFLEX, and the data were analyzed using CytExpert software. All staining was performed according to the manufacturer’s protocols. The control group without antibodies and staining was used for gating, and single-color stain controls were used to enable correct compensation.

### Small Interfering RNA (siRNA) and transfection

siRNA targeting CD74 or scramble sequences were purchased from Research Cloud Biology (China). siRNA targeting CD74 was transfected into macrophages derived from THP-1 with lipofectamine 3000 (Invitrogen) according to the manufacturer’s instructions. The sequences of siRNA CD74 are as follows:

1# /rC//rU//rC//rC//rC//rA//rA//rG//rC//rC//rU//rG//rU//rG//rA//rG//rC//rA//rA/TT; /rU//rU//rG//rC//rU//rC//rA//rC/lr A//rG//rG//rC//rU//rU//rG//rG/irG//rA//rG/TT 2# /rC//rC//rG//rC//rC//rU//rA//rC//rU//rU//rC//rC//rU//rG//rUllrA /rC//rC/lrA/TT; /rU//rG/irG//rU//rA//rC//rA//rG//rG//rA//rA//rG//rU//rA//rG/rG//rC//rG//rG/TT

### RNA extraction and qRT-PCR

TRIzol (LIFE Ambion, USA) was used to extract the total RNA of tissues and cells, and the concentration was determined. Total RNA was reverse transcribed into cDNA using the HiScript III RT SuperMix for qPCR (+gDNA wiper) (Vazyme, R223-01). Then, we amplified the aimed gene fragment and detected the relative expression with the SYBR Green qPCR kit (TOYOBO, Japan). Quantitative real-time PCR was performed for 40 cycles. All experiments were conducted at least three times. Relative gene expression levels were calculated by the 2-CT method. Primer sequences for detecting human CD74 mRNA expression are 5’-GACGAGAACGGCAACTATCTG-3’ and 5’-GTTGGGGAAGACACACCAGC-3’.

### Phagocytosis assay

Macrophages derived from THP-1 and CaSki cells were digested with Trypsin (Macgene) and stained with the Cell Plasma Membrane Staining Kit with DiI (Red Fluorescence) or DiO (Green Fluorescence) (Beyotime, Jiangsu, China), respectively. The macrophages were plated at a density of 5 × 10^4^ cells per well in a 24-well tissue-culture plate, and 2 × 10^5^ CaSki cells were used for staining per experiment. The cocultured cells were incubated in serum-free medium for another 3 h, then analyzed by flow cytometer, and imaged by a fluorescence microscope. The control group without antibodies and staining was used for gating, and single-color stain controls were used to enable correct compensation.

### Cell migration and proliferation assays

Standard transwell inserts (0.8 μm, Corning, USA) were used to detect cell migration. Cells were plated into the upper compartment with 500 μl of serum-free medium, and the lower compartment was filled with medium containing 10% fetal bovine serum. After 24 to 48 h incubation at 37 °C, cotton swabs were used to remove the noninvaded cells on the filter’s upper surface, and the cells were fixed for two minutes in 100% methanol. Invaded cells on the lower side of the filter were stained with 0.5% crystal violet for 20 min, and images were captured using a microscope.

The CCK-8 (Cell Counting Kit-8) assay kit (Beyotime, China) was used to evaluate the level of cell proliferation. Briefly, the 10 μl Cell Counting Kit solution was added to the culture medium and incubated for 3 h. The absorbance was determined at 450 nm wavelength with a reference wavelength of 630 nm.

## Acknowledgments

We thank the Translational Medicine Core Facility of Shandong University for the consultation and instrument availability that supported this work. We also thank the Model Animal Research Centre of Shandong University for mouse housing and care. This work was supported by the National Key Technology Research and Developmental Program of China (grant number No. 2022YFC2704401), the National Natural Science Foundation of China (grant numbers 82172940, 82103425, 82303055), the Shandong Provincial Natural Science Foundation (grant numbers ZR2021QH187, ZR2021QH044, ZR2023QH312), and the Jinan City “20 New University” independent innovation group (grant number 2021GXRC027).

## Competing interests

The authors declare that they have no competing interests.

## Data and materials availability

The raw scRNA-seq data reported in this paper have been deposited in the Genome Sequence Archive^61^ in National Genomics Data Center^62^, China National Center for Bioinformation / Beijing Institute of Genomics, Chinese Academy of Sciences (GSA-Human: HRA005959) under project PRJCA021022 that are publicly accessible at https://ngdc.cncb.ac.cn/gsa-human.

## Notes

### Competing Interest Statement

The authors have declared no competing interest.

