## Supplement Figures for "Targeting CD74-positive macrophages improves neoadjuvant therapy in cervical cancer as revealed by single-cell transcriptomics analysis"

#### Supplementary Information

##### Supplementary Figure 1

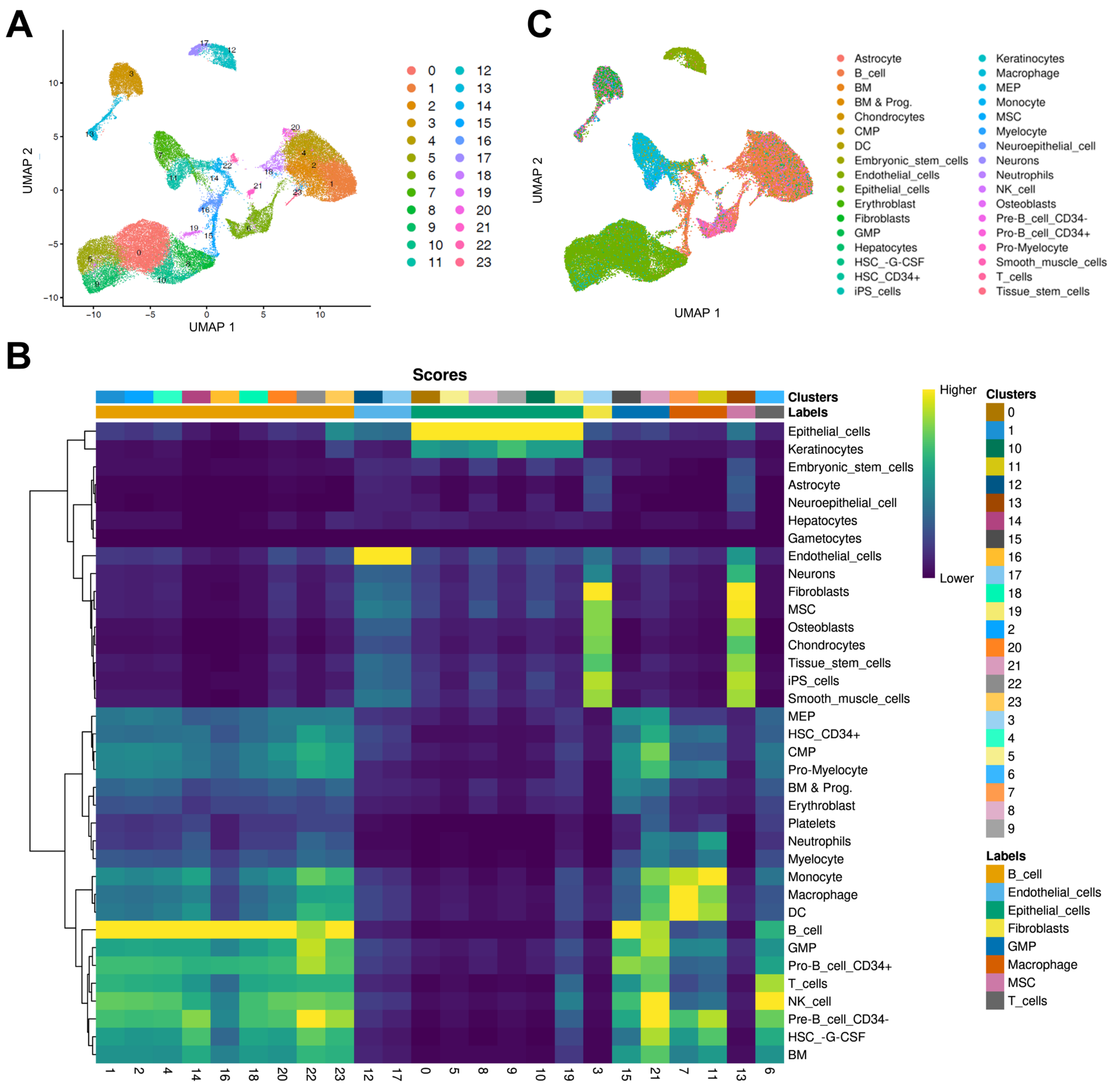


**Supplementary Figure 1. Cell type annotation of scRNA-seq data.** (A) UMAP projection of 24 raw clusters after dimension reduction. (B) The singleR score of the 36 cell types was calculated in the 24 clusters. The potential cell types were labeled in 8 colors by SingleR. (C) Initial single cell level annotation by SingleR. These results provide evidence for manual correction.

**Supplementary Figure 2**

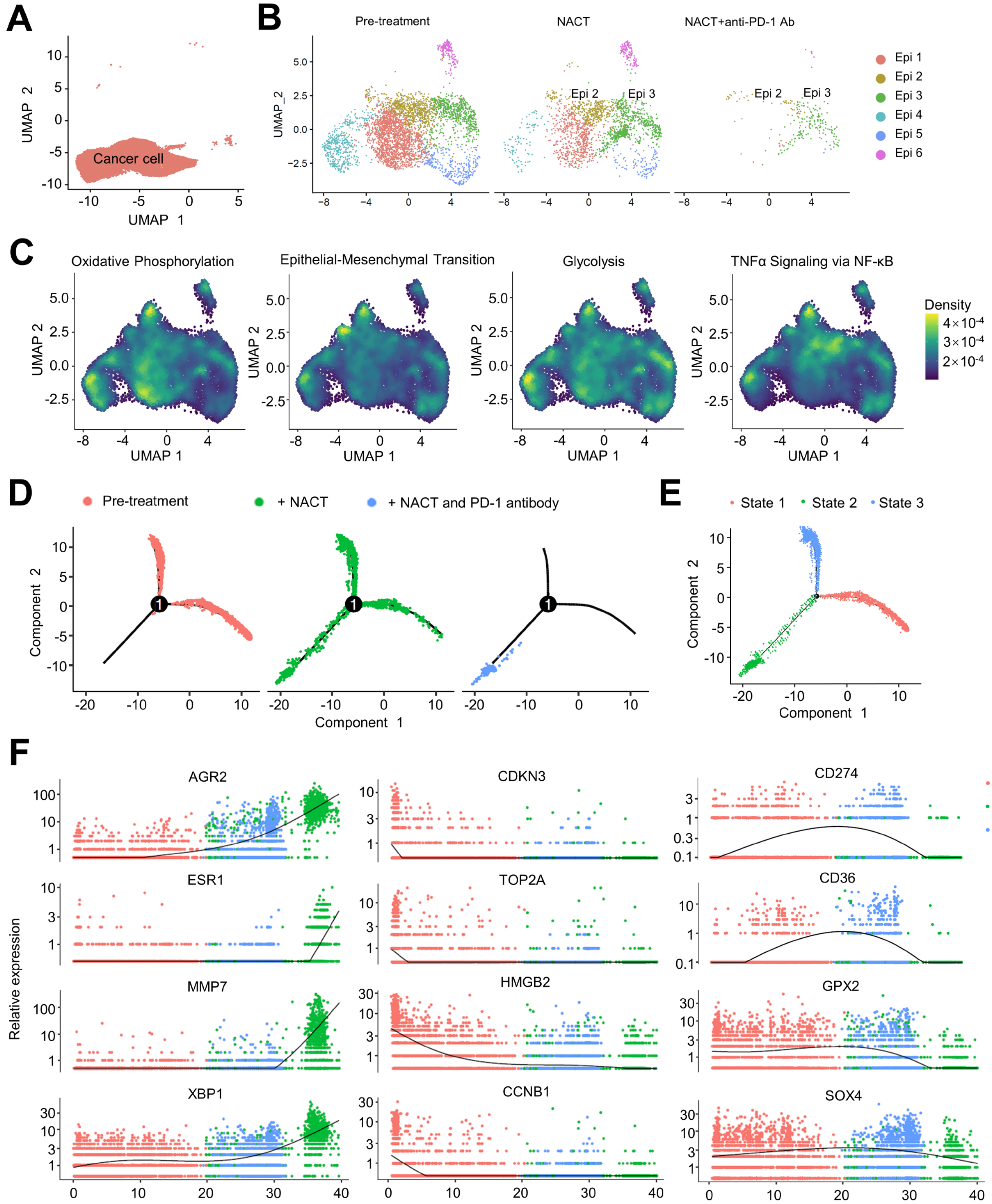


**Supplementary Figure 2. Enriched pathways of epithelial cancer cell subclusters and pseudo-time analysis.** (A) The cancer cell population before reducing the dimension. (B) UMAP projection of the six epithelial subgroups colored by different stages of NACT combined anti-PD-1 Ab therapy. (C) UMAP projection of oxidative phosphorylation, epithelial-mesenchymal transition, glycolysis, and TNFα signaling via NF-κB pathways density. (D-E) The unsupervised transcriptional trajectory of epithelial cells from Monocle (version 2), colored by different stages of NACT combined anti-PD-1 Ab therapy (D) and different transcriptional states (E). (F) The dynamic expression level of marker genes changed along with pseudo-time colored by different transcriptional states.

### Supplementary Figure 3

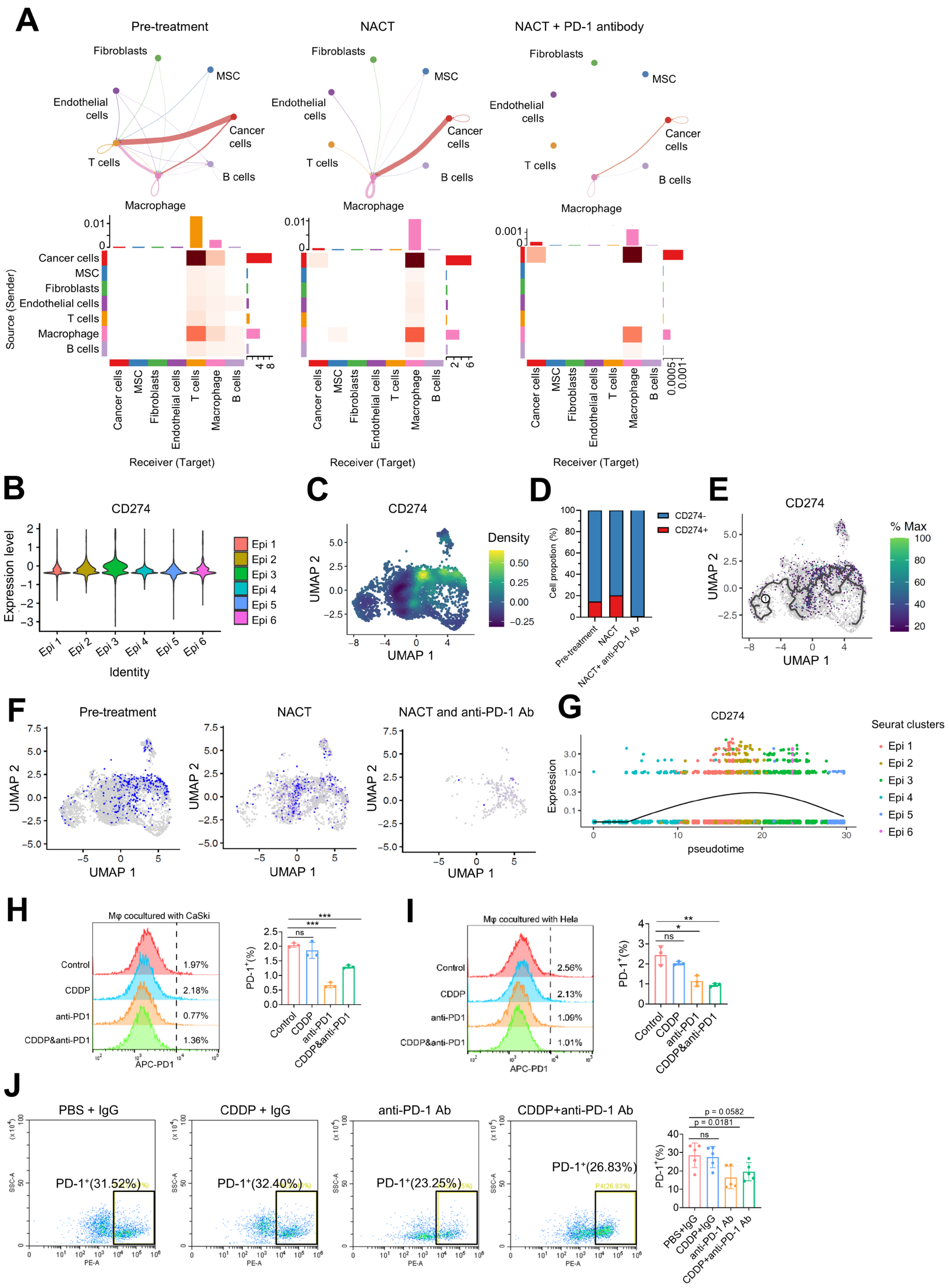

**Supplementary Figure 3. Cell communication between seven cell types and the effect of combined therapy on CD274 (PD-L1) and PD-1 expression.** (A) The interaction strength of significant ligand-receptor pairs between any pair of two cell populations. The edge width is proportional to the normalized interaction strength of ligand-receptor pairs. (B) Violin plot of CD274 (PD-L1) expression level in 6 epithelial subgroups. (C) UMAP projection of CD274 expression density in epithelial cells. (D) Proportion of CD274 positive and negative cells in pre-treatment, NACT, NACT + anti-PD-1 Ab groups. (E) Pseudo-time trajectory of epithelial cells and CD274 normalized expression. (F) UMAP projection of CD274 normalized expression in pre-treatment, NACT, NACT + anti-PD-1 Ab groups. (G) CD274 expression changes along with pseudo-time. Colored by epithelial subgroups. (H, I) PD1 positive cell percentage was determined by flow cytometry of macrophages cocultured with CaSki (H) and HeLa (I) cervical cancer cells in the control, CDDP, anti-PD-1, and CDDP combined with anti-PD-1 Ab group. (J) The PD-1 expression of macrophages isolated from the subcutaneous tumor microenvironment of the mice model was detected by flow cytometry. The p-value was obtained by a two-tailed unpaired Student's t-test, and the results are presented as the mean ± SD. $*p < 0.05$, $**p < 0.01$, $***p < 0.001$

**Supplementary Figure 4**

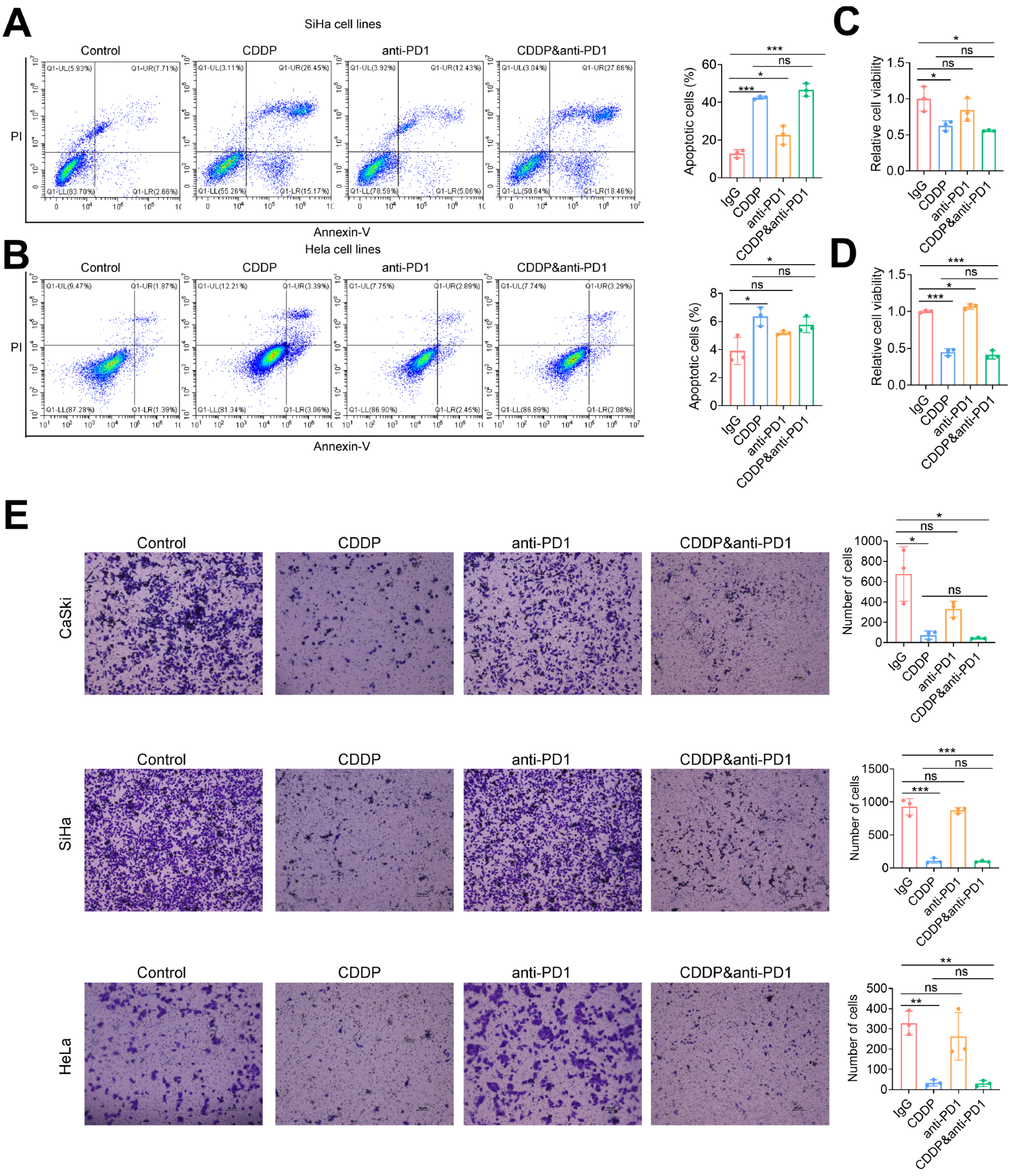


**Supplementary Figure 4. Anti-PD-1 Ab combination treatment did not affect cervical cancer cell viability, apoptosis, and migration ability without macrophage co-culture.** (A, B) Apoptotic cell percentages of HeLa (A) and SiHa (B) cells treated with anti-PD-1 Ab and/or CDDP. The sum of the upper right and lower right quadrants shows the percentage of apoptotic cells in the total measured cell population. Statistical graphs and representative experiments were shown. (C, D) The proliferative ability of HeLa (C) and SiHa (D) cells. (E) The migration ability of the cervical cancer cells treated with anti-PD-1 Ab and/or CDDP was determined by transwell assay. The number of cells stained in purple indicates the level of migration ability. The p-value was obtained by a two-tailed unpaired Student's t-test, and the results are presented as the mean ± SD. *p<0.05, **p < 0.01, ***p<0.001

#### Supplementary Figure 5

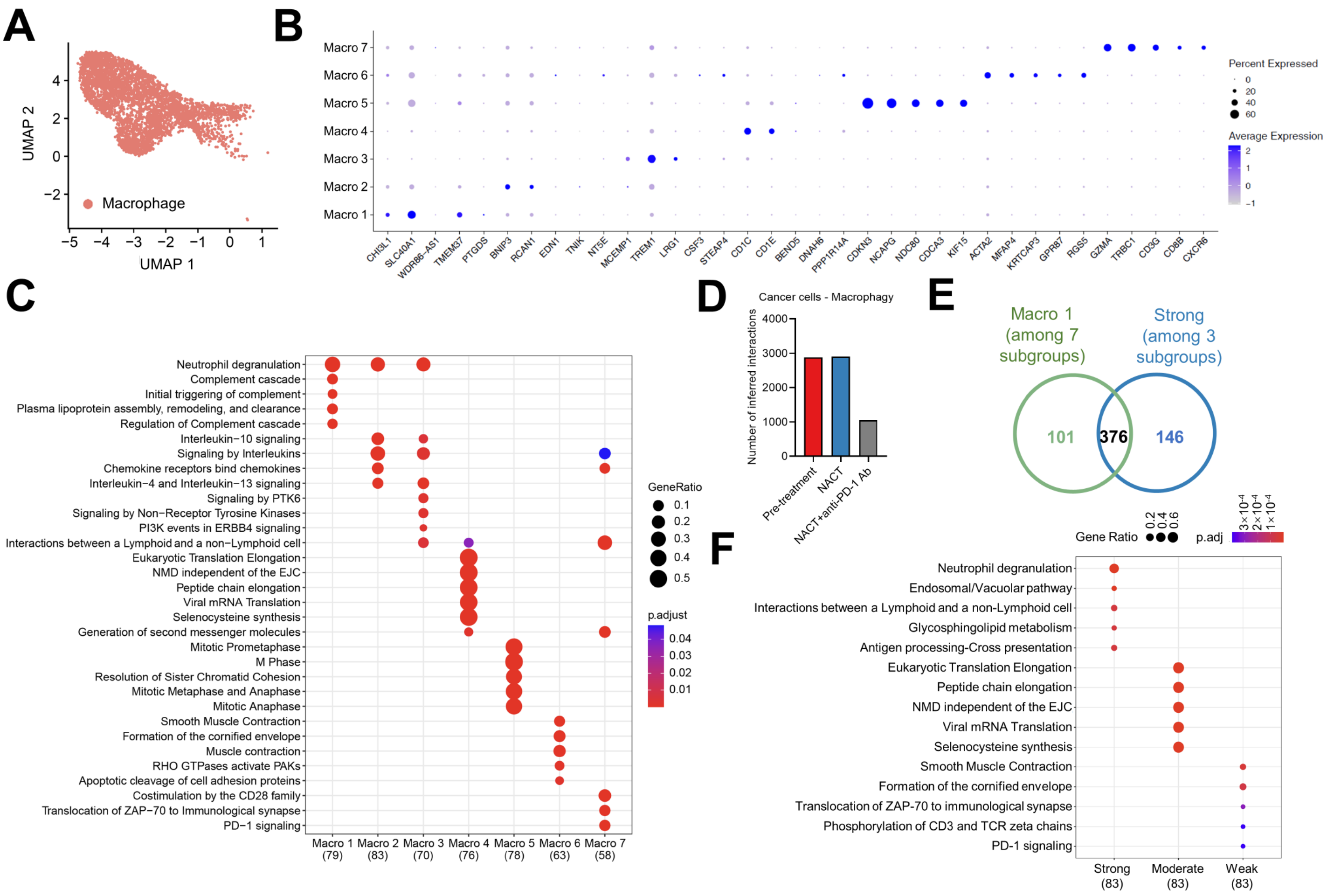


**Supplementary Figure 5. Marker genes and enriched pathways of macrophage subclusters.** (A) UMAP projection of 7 subgroups generated from sub-clustering macrophages. (B) Dot plot of average expression and expression percentage of marker genes for seven macrophage subgroups. (C) Dot plot of the enriched Reactome pathways of the seven macrophage subgroups. (D) The number of inferred interactions among cancer cell subclusters and macrophage subclusters before and after NACT treatment and the anti-PD-1 Ab combination. (E) Venn diagram of the genes shared by the Macro 1 and Strong subgroups. (F) Dot plot of the enriched Reactome pathways of the Strong, Moderate, and Weak macrophage subgroups. The dot size is proportional to the gene ratio and the color indicates the adjusted p-value.

**Supplementary Figure 6**

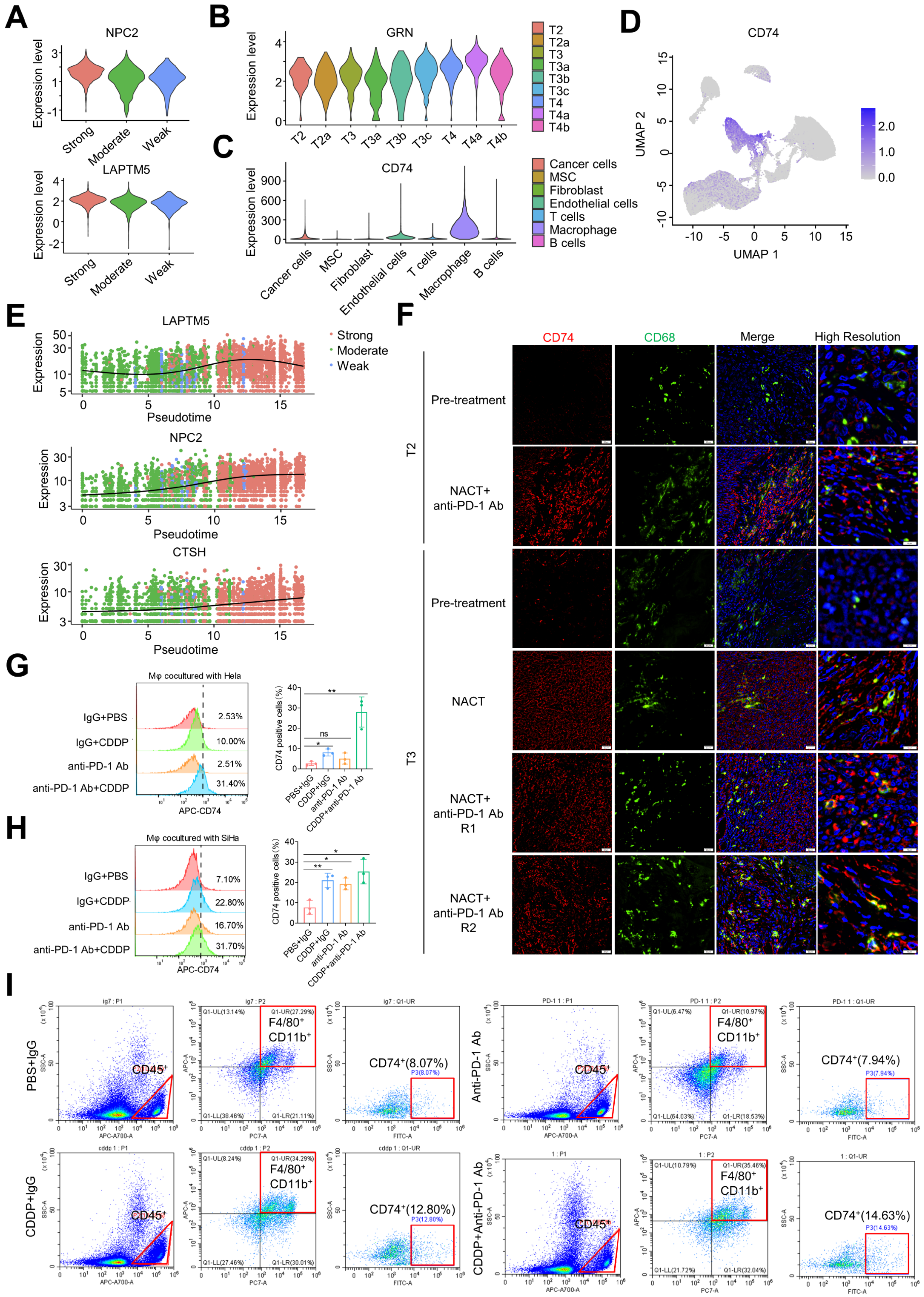

**Supplementary Figure 6. The dynamic expression changes of candidates participate in macrophage-cancer cell communication.** (A) Violin plot of the NPC2 and LAPTM5 expression in the Strong, Moderate, and Weak subgroups. (B) The normalized GRN expression level after pre-treatment, NACT, and anti-PD-1 Ab combination therapy (NACT+anti-PD-1 Ab) in T2, T3, and T4 patients. (C) The normalized CD74 expression level in seven cell types. (D) The UMAP projection of CD74 normalized expression. (E) The LAPTM5, NPC2, and CTSH expression dynamic change along with the pseudo-time, colored by the Strong, Moderate, and Weak subgroups. (F) The immunofluorescence assay showed the CD74 (red) and CD68 (green) expression in the tissues of patients T2 and T3 receiving NACT and anti-PD-1 Ab combination therapy. (G, H) SiHa (G) or HeLa (H) cells were cultured with macrophages derived from THP-1 cells. Anti-PD-1 Ab and CDDP were used to establish a model of NACT combined with immunotherapy in vitro. The percentage of CD74 positive macrophage population in total cells was detected by flow cytometry. (I) Gating strategy and representative images of flow cytometry. The percentage of CD74+ macrophages in single-cell suspensions isolated from mice subcutaneous tumor tissues. CD45 antibody is used to screen monocytes in cell suspension. In the CD45-positive cell population, the macrophages expressing both F4/80 and CD11b were gated. The p-value was obtained by a two-tailed unpaired Student's t-test, and the results are presented as the mean ± SD. *$p<0.05$, **$p < 0.01$, ***$p<0.001$

**Supplementary Figure 7**

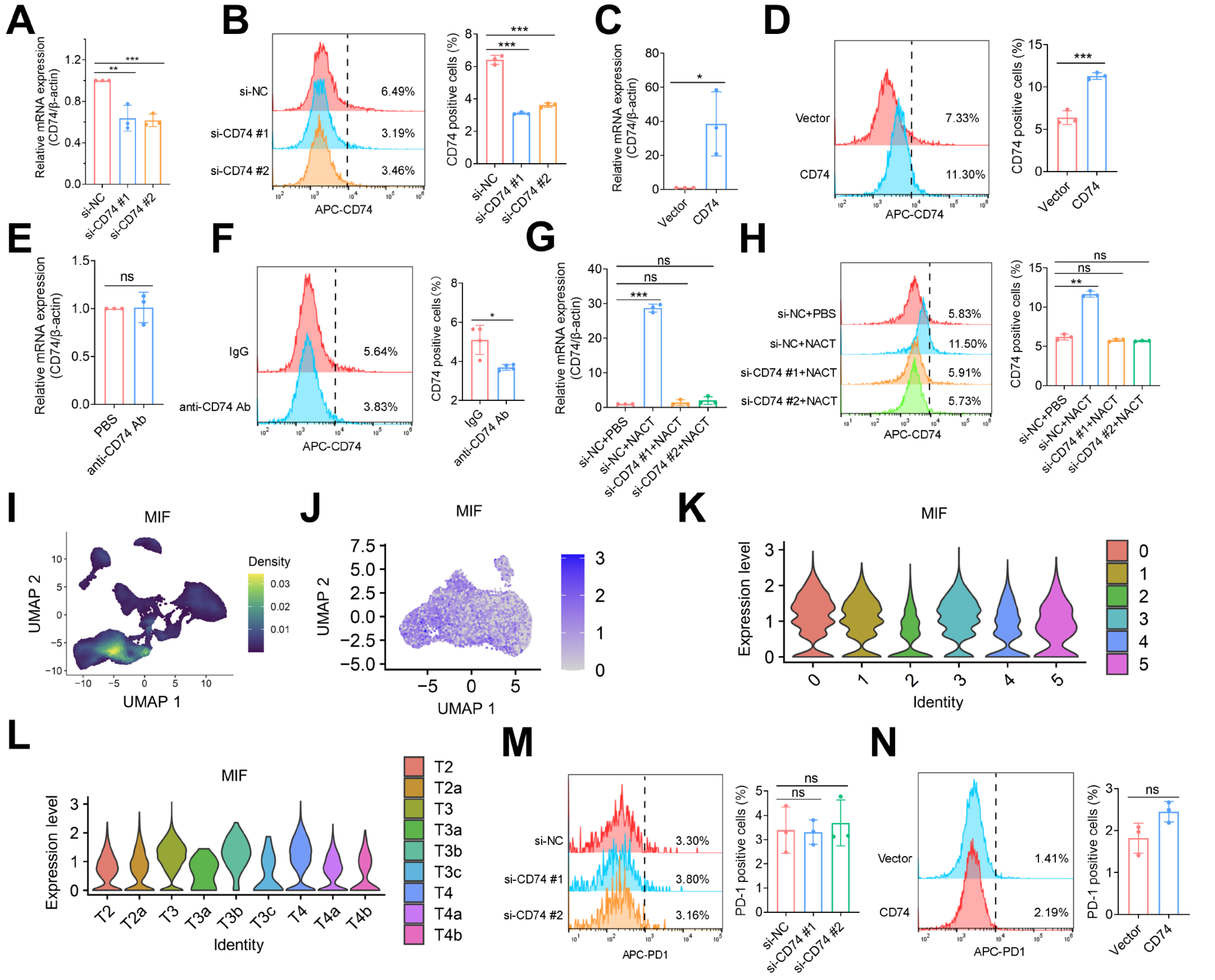


**Supplementary Figure 7. CD74 expression after anti-CD74 Ab, platinum treatment, and MIF expression in epithelial cell subgroups and different samples.** (A) The CD74 mRNA expression in THP-1-derived macrophages after siCD74 or siNC transfection. (B) The proportion of CD74 positive cells detected by flow cytometry after siCD74 or siNC transfection (C) The CD74 mRNA expression in THP-1-derived macrophages transfected with CD74 overexpression plasmid. (D) The proportion of CD74 positive cells after transfected with CD74 overexpression plasmid. (E) The CD74 mRNA expression in macrophages after treated with anti-CD74 Ab. (F) The proportion of CD74 positive cells after anti-CD74 Ab treatment. (G) The mRNA expression of CD74 was knocked down in THP-1-derived macrophages stimulated by CDDP. (H) The CD74 positive macrophage percentage was detected by flow cytometry. (I, J) UMAP projection of MIF normalized expression in all types of cells (I) and epithelial cell subgroups (J). (K, L) Violin plot of MIF expression in six epithelial cell subgroups (K) and 9 tissues (L). (M, N) PD-1 expression was determined by flow cytometry after CD74 knockdown (M) and overexpression (N). (M) The proportion of PD-1 positive cells after transfected with siCD74 or siNC. (N) The proportion of PD-1 positive cells after transfected with CD74 overexpression plasmid. The p-value was obtained by a two-tailed unpaired Student's t-test, and the results are presented as the mean ± SD. *p<0.05, **p < 0.01, ***p<0.001
